# Global genetic heterogeneity in adaptive traits

**DOI:** 10.1101/2021.02.26.433043

**Authors:** William Andres Lopez-Arboleda, Stephan Reinert, Magnus Nordborg, Arthur Korte

## Abstract

Understanding the genetic architecture of complex traits is a major objective in biology. The standard approach for doing so is genome-wide association studies (GWAS), which aim to identify genetic polymorphisms responsible for variation in traits of interest. In human genetics, consistency across studies is commonly used as an indicator of reliability. However, if traits are involved in adaptation to the local environment, we do not necessarily expect reproducibility. On the contrary, results may depend on where you sample, and sampling across a wide range of environments may decrease the power of GWAS because of increased genetic heterogeneity. In this study, we examine how sampling affects GWAS for a variety of phenotypes in the model plant species *Arabididopsis thaliana*. We show that traits like flowering time are indeed influenced by distinct genetic effects in local populations. Furthermore, using gene expression as a molecular phenotype, we show that some genes are globally affected by shared variants, while others are affected by variants specific to subpopulations. Remarkably, the former are essentially all *cis*-regulated, whereas the latter are predominately affected by *trans*-acting variants. Our result illustrate that conclusions about genetic architecture can be incredibly sensitive to sampling and population structure.

## 1 Introduction

Genome-wide associations studies (GWAS) have become the standard tool for analysing the relationship between genotype and phenotype in populations. Pioneered in human genetics (Hirschhorn and Daly 2005), GWAS are now widely used in many different species to infer trait architecture and identify causal variants. Much less work has been done on comparing architectures across populations. Repetition of GWAS in different human populations or samples have mostly been used in meta-studies to improve power, although awareness is growing that genetic architecture may be different between populations (Martin et al. 2017; Sohail et al. 2019; Berg et al. 2019). However, when working on traits that are likely to be involved in local adaptation, there is every reason to expect differences in the underlying genetic architecture. How does this genetic heterogeneity affect GWAS and what can we learn from it? To address these questions, we used data from the model plant species *Arabidopsis thaliana*, which occurs throughout the northern hemisphere, and has been shown to be locally adapted (Fournier-Level et al. 2011; Hancock et al. 2011; Ferrero-Serrano and Assmann 2019). Our general strategy was to compare the GWAS results from a global sample to various regional subsamples. We started using flowering time as a trait, since it is well studied, subject to strong selection (Flowers et al. 2009, Ågren et al. 2017) and well-understood molecularly in *Arabidopsis thaliana* (Henderson and Dean 2004) and in other plant species (Weller and Ortega 2015). We also analysed stomata size and cauline leaf number as additional examples of “real” phenotypes, and compared the results with simulations to establish how GWAS in subpopulations would be expected to behave under simple models. Finally, we performed GWAS on gene expression levels to investigate whether gene regulation shows evidence of local adaptation.

## 2 Analysis

### 2.1 Flowering time is affected by different alleles in different populations

We used publicly available flowering data, measured in growth chambers at 10°C for over 1,000 accessions (The 1001 Genomes Consortium 2016). We restricted our analysis to 888 accessions from Europe and divided those into eight semi-arbitrary subpopulations of approximately equal sizes using only geographic information: Southern Iberian Peninsula (SIP), Northern Iberian Peninsula (NIP), Germany, France/UK, Central Europe, Skåne (the southernmost province of Sweden), Northern Sweden (Sweden excluding Skåne), and Eastern Europe (Fig. 1A, Table S1). All subpopulations had highly variable flowering times, with only the two Swedish ones being generally later-flowering (Fig. 1B).

**Fig 1:**
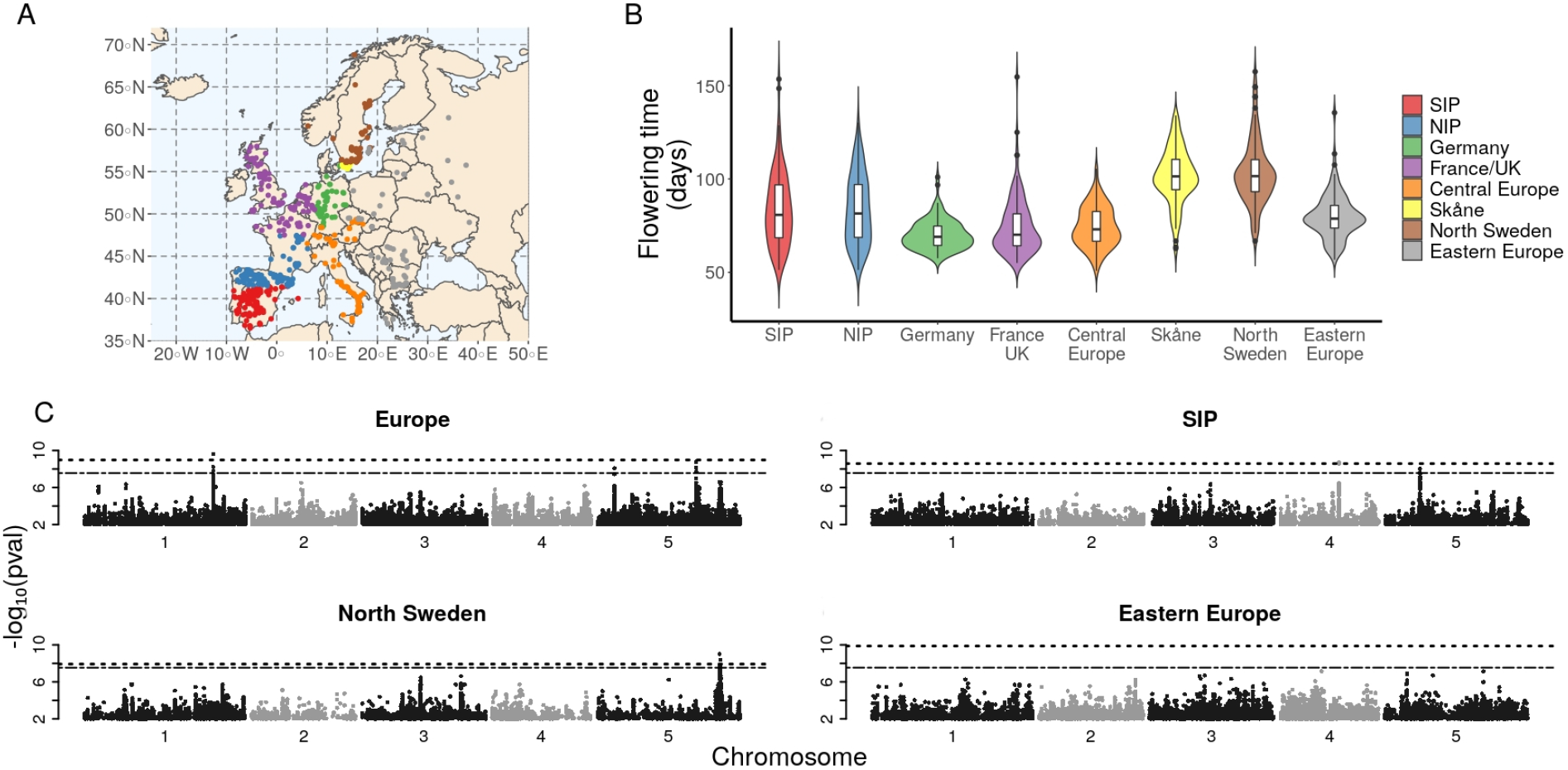
GWAS of flowering time across Europe. (**A**) Origin of the 888 European *A. thaliana* accessions, with eight designated subpopulations in different colors. (**B**) The distribution of flowering time in the subpopulations. (**C**) Manhattan plots of GWAS results for the whole European populations and three of the eight subpopulations. Dashed and dash-dotted lines indicate Bonferroni- and permutation-based 5% significance thresholds, respectively.

We performed GWAS on the entire European population as well as in the different subpopulations. Note that, although the subpopulations are small (*n* = 103–119), flowering time has extremely high heritability and major polymorphisms are believed to be common (Mouradov et al. 2002). Simulations suggested that power should be sufficient to identify such polymorphisms (Table S1), and this is born out by results that pinpoint several well-known genes (Figs. 1C and S1, Table 1).

**Table 1:**
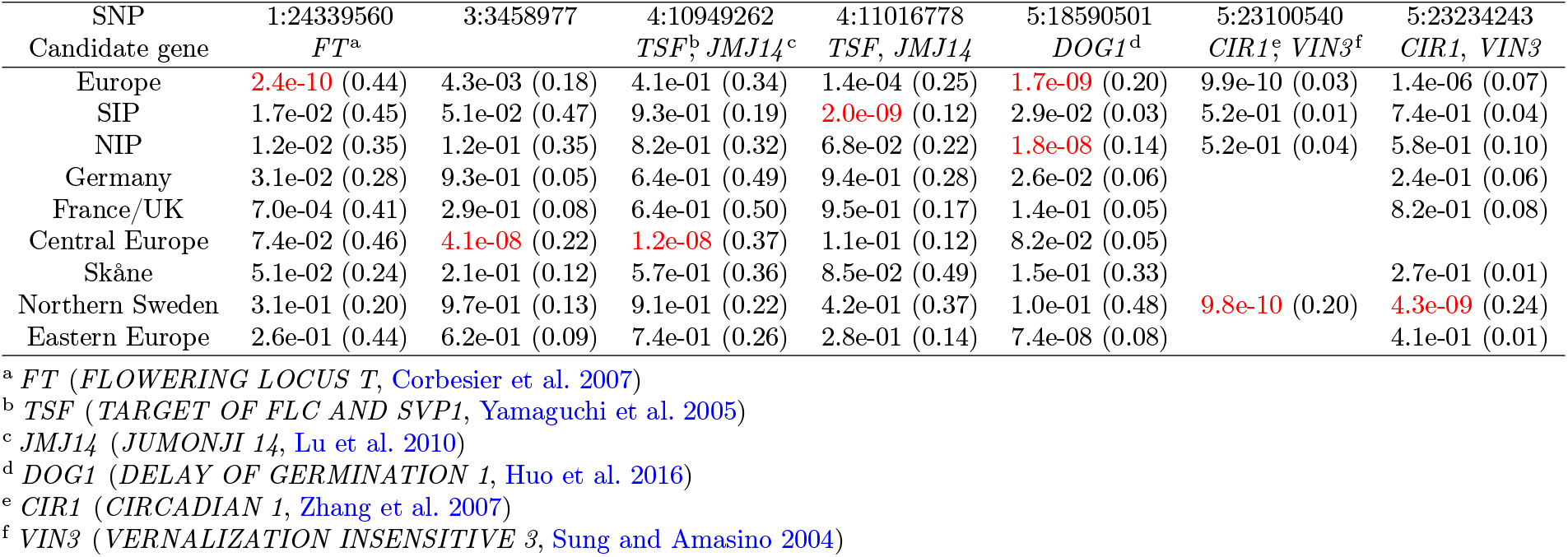
Significant SNPs (chromosome:position) in the GWAS of different subpopulations. Entries are “*p*-value (minor allele frequency)”, with genome-wide significance using a 5%-permutation-based threshold shown in red. Candidate genes were assigned to the SNPs from a list of 306 flowering time genes (Bouché et al. 2016) using 10 kb window.

Using a permutation-based threshold (Freudenthal et al. 2019), we identified genome-wide significant associations in four of the subpopulations as well as in the full European population (Tables 1 and S2–S3). The results differed strikingly, with only one association, near *DOG1*, showing any signs of significance in more than one subpopulation. This association was significant in NIP, almost significant in Eastern Europe, and also significant in the full population (Fig. S2). *DOG1* is an extensively studied gene involved in the regulation of seed dormancy (Huo et al. 2016; Kerdaffrec et al. 2016), but has also been identified in GWAS for flowering time (Atwell et al. 2010).

Whether a causative polymorphism is detected or not depends on its effect size, its frequency, and whether it is “tagged” by a marker included in the study. The latter is a major concern when comparing human populations, because sparse SNP data are used, and patterns of linkage disequilibrium can differ greatly between populations (Martin et al. 2017). While this explanation cannot be excluded here, it is likely to be much less important, because we are using dense SNP data from whole-genome re-sequencing. Compared to a standard human GWAS, we are using four times as many markers in a genome that is 25 times smaller, but in which linkage disequilibrium is roughly as extensive (Nordborg et al. 2002; The 1001 Genomes Consortium 2016).

The other two explanations are more interesting. For polymorphisms involved in local adaptation, allele frequencies are expected to differ between geographic regions, and the *VIN3* association (5:23100540) may be an example of this. The minor allele at this locus appears to be associated with late flowering across Europe, but is too rare to be detected except in Northern Sweden (Table 1 and Fig. S3).

However, we also see several examples of SNPs that are common everywhere, but show no sign of being associated with flowering time except for a single population. As noted above, differences in linkage disequilibrium with closely linked unobserved causal polymorphisms are impossible to rule out, but we think it is more likely that the difference is the broader genetic background. The genetic background could influence the effect size through epistatic interactions with other loci or via genome-wide linkage disequilibrium caused by population structure or selection (Yu et al. 2006; Vilhjálmsson and Nordborg 2012). In our analyses, the effect of the genetic background is estimated using a mixed model, and the single-locus association is based on the Best Linear Unbiased Predictor (BLUP) of the marginal effect size (Yu et al. 2006). As the name indicates, this estimate should be unbiased, but there is no guarantee that this will be the case if the assumptions of the model (notably a polygenic, additive background) are violated.

To confirm that these conclusions are not limited to genome-wide significant SNPs, we next compared all SNPs with *p <* 10^*−*4^. In agreement with the results just presented, only eight of over 5,000 sub-significant SNPs were shared among subpopulations, and associations were never shared among more than two (Fig. 2A and Table S4). Note that this are far more associations than expected by chance; indeed, even the overlap is higher than expected (Fig. S4A). Also notable is that shared sub-significant associations are clearly clustered in regions that tend to be common between subpopulations (Fig. 2B) and significantly fewer shared regions are detected in the simulations (Fig. S4B and Fig. S5). Although regions shared among multiple subpopulations are clearly associated with known flowering time genes (Table S5 and File S1), no significant enrichment had been observed.

**Fig 2:**
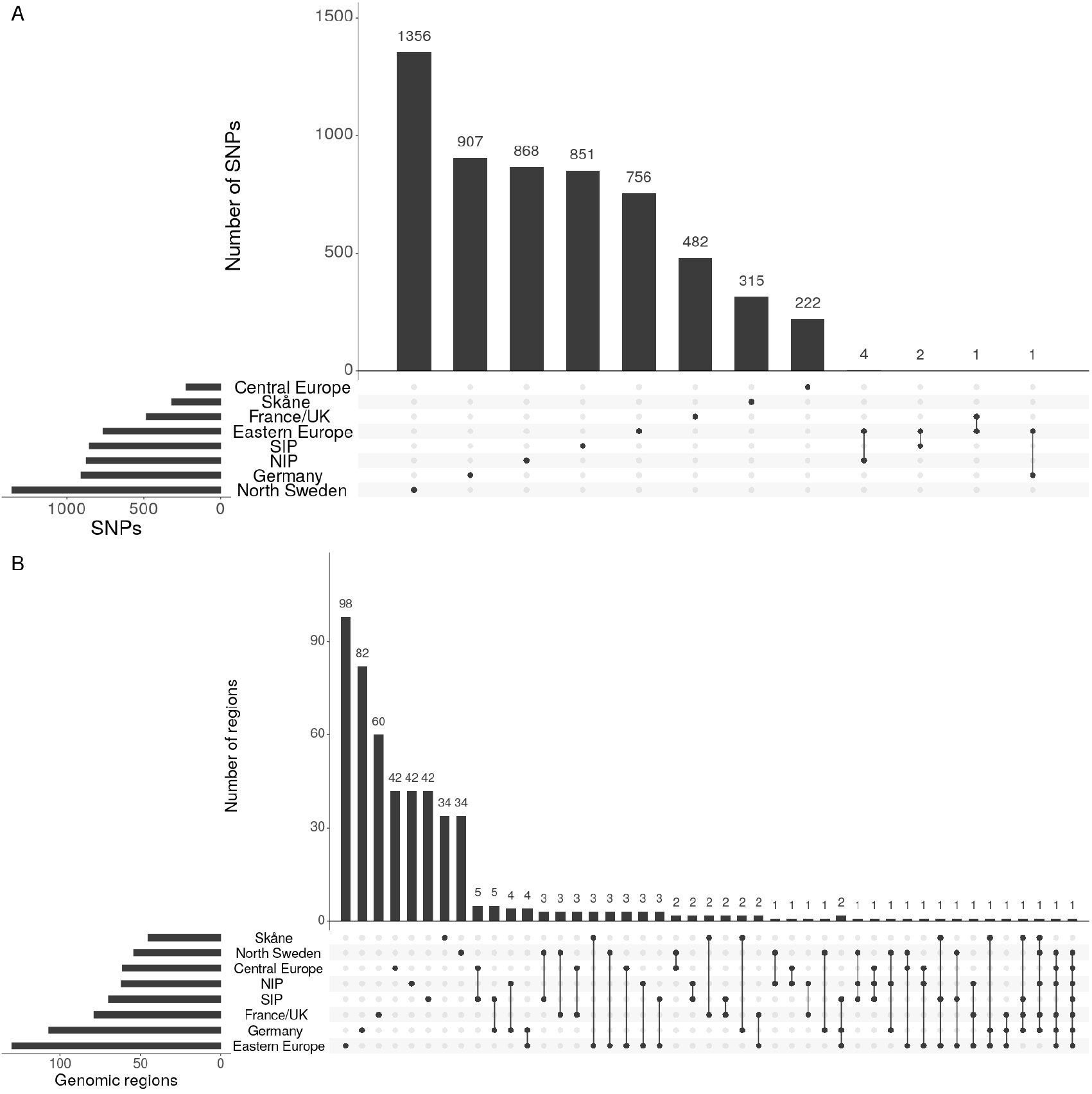
Sharing of sub-significant (*p <* 10^*−*4^) associations. (**A**) Histogram of the number of associated SNPs in each subpopulation and shared between subpopulations. (**B**) Histogram of the number of associated genomic regions in each subpopulation and shared between subpopulations.

There are two possible explanations why different SNPs pinpoint the same genomic region. The first is that the causal polymorphisms are absent from the data, and that different SNPs “tag” the (shared) causal polymorphisms in the different subpopulations. As noted above, given the high SNP density used, we do not believe this is a general explanation. More likely is extensive allelic heterogeneity, a phenomenon consistent with local adaptation, and well-demonstrated in *A. thaliana* (Atwell et al. 2010; Li et al. 2014; Kerdaffrec et al. 2016; Zhang and Jiménez-Gómez 2020).

To further investigate the putative heterogeneity, we estimated the polygenic overlap between subpopulations using a method that estimates the correlation of marker effects across different samples and thus tries to circumvent the problem of detecting causal variants (Frei et al. 2019). As anticipated, estimates of marker-effect correlations were low in all comparisons. As a contrast, the marker-effect correlation between flowering time at 10°C and 16°C (FT16) was significantly higher (Fig. S6).

### 2.2 Simulations suggest that local genetic architecture is detectable

Flowering time is the quintessential locally adaptive trait. It is difficult to know how unusual it is, because few traits have been measured in different populations in wild species. We looked at two publicly available data sets: stomata size and cauline leaf number, measured in 131 accessions from Sweden and 109 from the Iberian Peninsula (Fig. S7). However, the analysis was uninformative, as no genome-wide significant associations were identified (Fig. S8), and no overlap was found for sub-significant associations either (Fig. S9). Indeed, despite both phenotypes having high heritabilities (27-85%; Table S6), the joint p-value distribution was indistinguishable from noise, suggesting that GWAS is under-powered to detect causal alleles for these phenotypes, presumably because they are highly polygenic, and major alleles do not exist.

To gain insight into the power to detect causal alleles in this setting, we turned to simulations. Briefly, we simulated phenotypes using the 240 accessions used for the analysis of stomata size and cauline leaf number. A single randomly picked polymorphism was assumed to explain a fixed percentage of the phenotypic variation, in either the Swedish, the Iberian, or the merged population (see Methods for details). We calculated the power to identify the causal polymorphism as well as the number of false positive associations using a Bonferroni-threshold and a thousand simulations for each scenario (Tables S7–S8, Fig. S10). In summary, the simulations suggested that GWAS in our small populations have sufficient power to identify major alleles and population-specific effects — supporting our claim of local adaptation for flowering time, and also that major alleles for stomata size and cauline leaf number do not exist.

### 2.3 Gene expression can be regulated globally or locally

Finally, we carried out GWAS for gene expression data, available for a large sample of world-wide accessions (Kawakatsu et al. 2016). It seemed *a priori* likely that some genes are under local selection, while others are not. For comparison with the results above, and because reasonably dense local samples were available, we focused on 91 accessions from the Iberian Peninsula (IP) and 74 accessions from Scandinavia (termed SW, as nearly all accessions are from Sweden). We carried out GWAS for each subpopulation, as well as for the merged set of 165 accessions. Because of the small sample sizes, we only considered genes with high estimated heritability and for which simulations indicate sufficient power in all three populations (see Methods). These criteria led to the retention of 2,237 genes, 9% of the total (Table S9). We also excluded genes where inflated significance levels where observed: this further reduced the number of genes to 1,982.

Perhaps not surprisingly, 780 (39%) of these filtered genes revealed a genome-wide significant association (using a multiple-testing corrected threshold of *p <* 10^*−*10^) in at least one of the two subpopulations (typical results are shown in Fig. 3). These genes were divided according to the pattern of associations within and between subpopulations, with the intent to identify those with clear evidence for global vs. local genetic architecture (see Fig. S11 and Methods for details).

**Fig 3:**
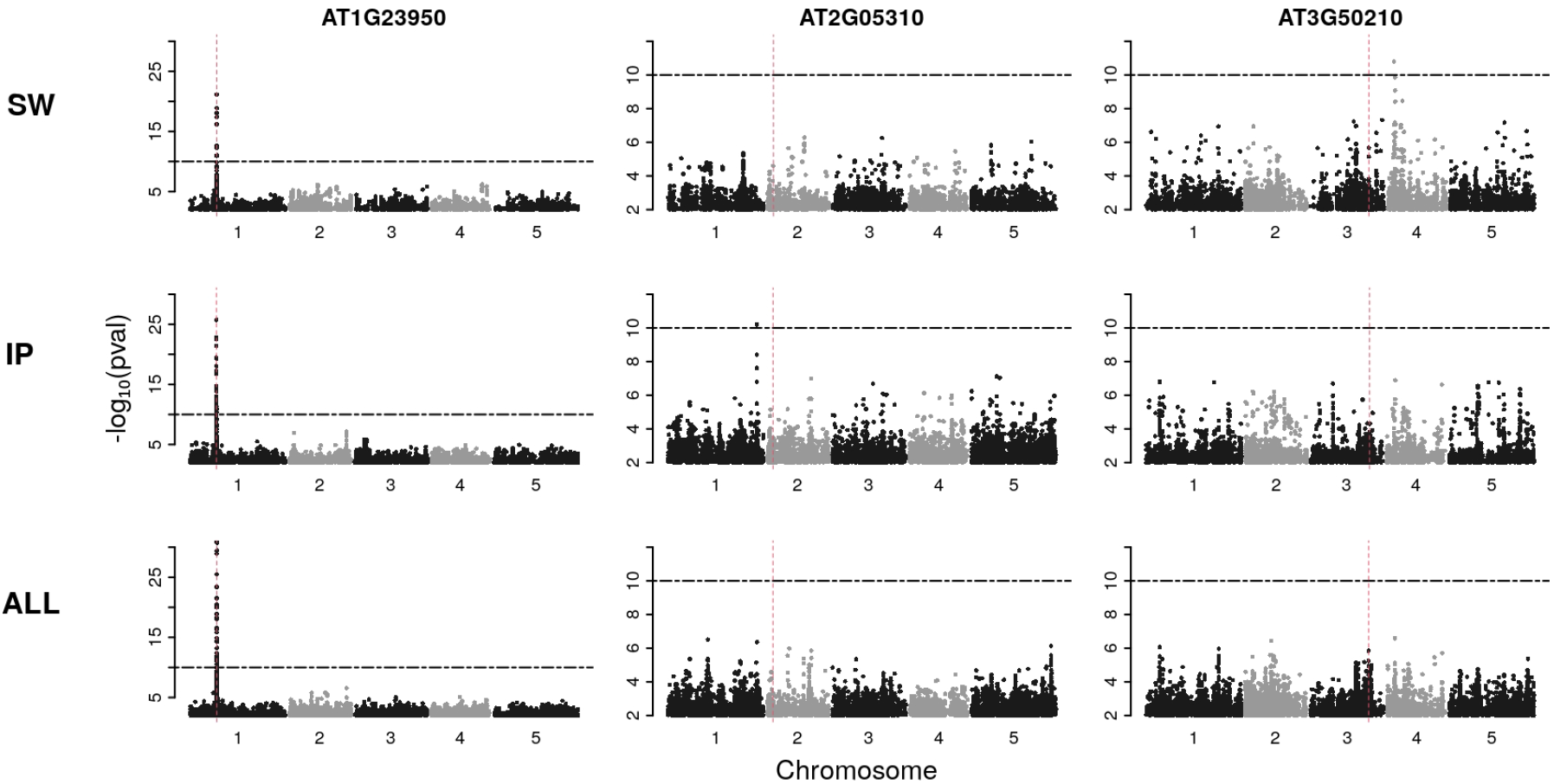
Manhattan plots from GWAS of expression levels for three different genes. The Columns show the results from genes representing different scenarios. The rows display the GWAS results of the analysis in the two subpopulations (SW and IP, respectively), or in the merged population (ALL). Horizontal dash-dotted lines indicate the significance threshold of *p <* 10^*−*10^. Vertical dashed lines show the position of the gene whose expression is being used as a molecular phenotype.

We found clear examples of both. Of the 780 genes with a significant association, 110 (14%) were significantly associated with the same SNP in both subpopulations (shared architecture), 25 (3%) were significantly associated with different SNPs in the same 50 kb genomic region in the both subpopulation (presumably allelic heterogeneity), 92 (12%) were significantly associated with different SNPs at distinct genetic regions in the two subpopulations (genetic heterogeneity), and 182 (23%) appeared to show an specific association in one subpopulation only (also genetic heterogeneity). The remaining are more ambiguous (Fig. S11).

Unexpectedly, we also found an extremely strong pattern of *cis-*vs. *trans-*regulation. Of the 110 genes with shared association between subpopulations, 99% were *cis*-regulated, whereas the opposite was true for genes with different regulation in the subpopulations. Here, 75% of the 182 genes that appeared to show an specific association in one subpopulation only were *trans*-regulated (Figs. 4 and S11).

**Fig 4:**
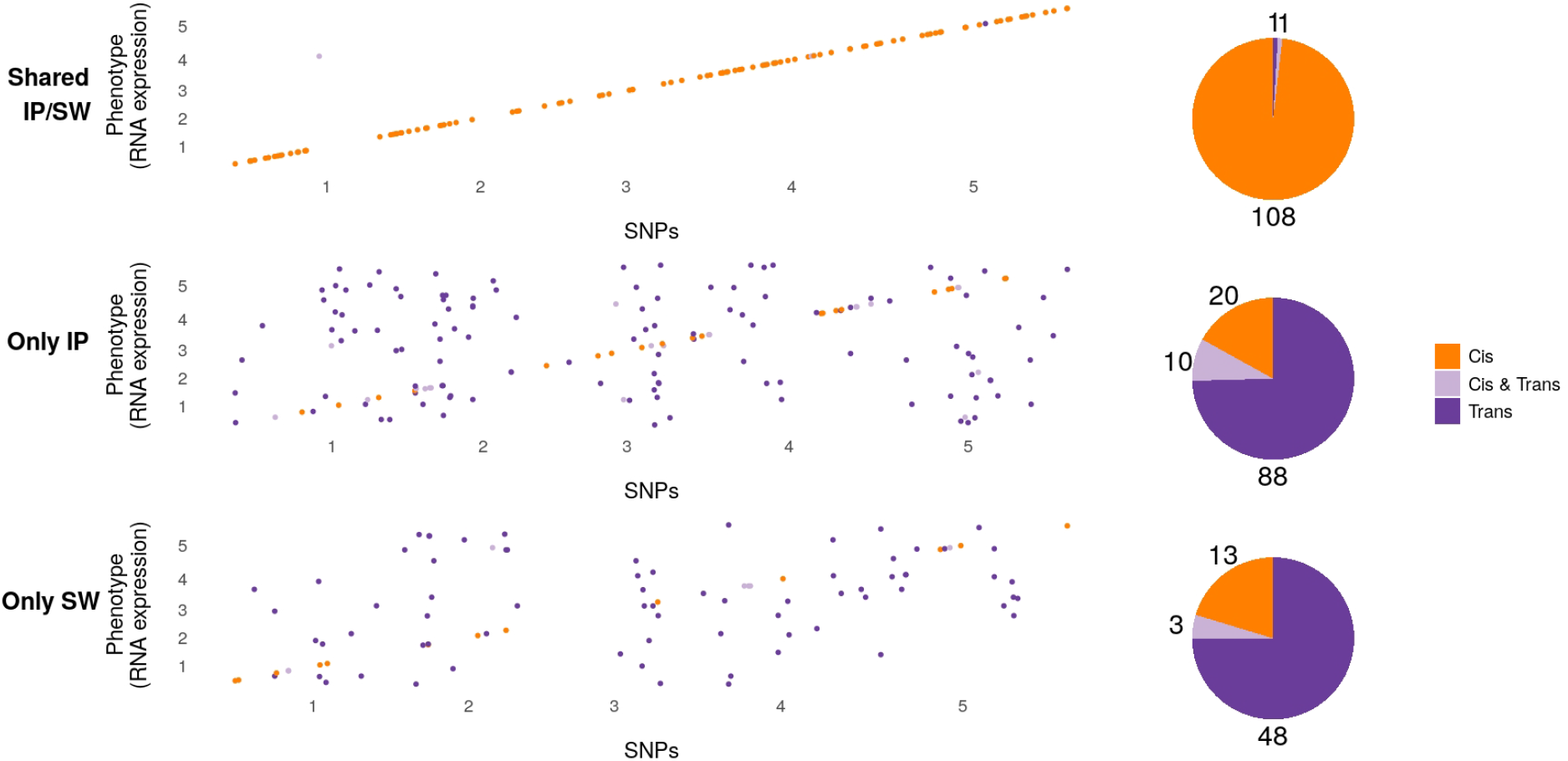
Summary of the difference between shared and non-shared GWAS results for expression data. Plots show the position (x-axis) of significant associations for each expressed gene (y-axis). Associations shared between subpopulations (top panel) are almost all in *cis*, whereas associations specific to one subpopulation (bottom panels) are mostly found in *trans*. Pie-charts show the number of genes in each category.

To confirm that these results reflect real differences between the subpopulations, we generated two random populations of the same size by randomizing the subpopulation labels. As expected, this recovered the shared *cis*-associations (157 genes showed shared associations, of which 94% are in *cis*, Figs. S14– S15). Non-shared associations were still mostly in *trans*, but there are less than half as many clearly subpopulation specific ones (Fig. S14–S16). This suggest that a substantial fraction of the specific associations found in the Scandinavian and Iberian populations are real. Further supporting this, only five genes showed a pattern of allelic heterogeneity in the analysis of the random, non-local subpopulations.

A GO-enrichment analysis found a significant enrichment for “ADP binding” among genes displaying the same global architecture, while no significant enrichment for those with specific local associations has been found. More anecdotally, the group of genes with shared variants contains many genes linked to primary metabolic pathways, as well as genes like *RPS5* (*RESISTANT TO P. SYRINGAE 5*), which is linked to bacterial and downy mildew resistance (Warren et al. 1998), and which is likely to be under global balancing selection (Tian et al. 2002). The set of genes, whose eGWAS results indicates local adaptation, contains for example genes related to flowering time regulation, like *AGL-20* (*AGAMOUS-LIKE 20* ; Lee 2000), and stress response, like *RCAR5/PYL11* (*REGULATORY COMPONENT OF ABA RECEPTOR 5/ PYRABACTIN RESISTANCE-LIKE 11* ; Lim and Lee 2020) and *HDA9* (*HISTONE DEACETYLASE 9* ; Zheng et al. 2016).

## 3 Discussion

It has been clear for over a decade that GWAS in plants often produce results that are strikingly different from those typically seen in humans. Major associations explaining substantial fractions of the phenotypic variance are found frequently, likely because this variance is adaptive, and the allelic variants are maintained by selection (Atwell et al. 2010; Huang et al. 2010). A prediction from this is that we do not necessarily expect GWAS results to replicate between populations, because many traits are likely to be involved in local adaptation. Here we use a simple analysis to show that this is very much the case for flowering time, a trait known to be important for local adaptation. We then show that the same is true for expression variation at many genes, and discover a striking pattern in that regulatory variants that are shared between populations are almost all in *cis*, whereas those that may be involved in local adaption are predominantly in *trans*.

That local adaptation would frequently involve *trans*-regulation is perhaps not surprising, as it seems likely that such adaptation generally involves expression changes at large number of loci, and this is surely easier to achieve using variation at upstream regulatory loci. It is also consistent with findings from *A. thaliana* that genotype-by-environment interactions in gene expression are mostly due to *trans*-acting variants (Clauw et al. 2016), and that, analogously, tissue-specific expression variation in humans also tends to be due to *trans*-acting variation (GTEx Consortium 2017).

The role of *cis*-regulatory variation under this scenario is less clear. Clauw *et al*. (2016), found that *cis*-regulatory variants had the same effect in different environments, suggesting that they could only be involved in local adaptation through highly polygenic allele-frequency changes. Alternatively, most of the *cis*-regulatory variations could simply be neutral. It should be noted, however, that we found many genes with allelic heterogeneity in their *cis*-regulation, suggesting that local adaptation via *cis*-regulation is possible as well.

Our findings also have important implications for the design and interpretation of GWAS. As part of the “1001 Arabidopsis Genomes Consortium”, we have often been asked “Which subset of accessions should I use?”. This paper shows that there is no simple answer. Clearly, what you find depends on where you look, and the optimal design depends on the question as well as on the phenotype. A global sample may not have the power to detect locally important allelic variation, and a local sample may not even contain globally important variants. Depending on the nature of reality, you will always miss some part of the picture, and if you are not aware of this, you may draw the wrong conclusions. For example, the relatively importance of *cis*-versus *trans*-regulation has been much debated (reviewed in Signor and Nuzhdin 2018), but this paper show that the answer likely depends on how you sample. In conclusion, GWAS works, but should be used with caution.

## 4 Methods

### 4.1 Plant material and phenotypic data

The phenotypic data used in this study, were obtained from the *A. thaliana* phenotype repository AraPheno (Seren et al. 2016). The genotypic data were obtained from the 1001 Genomes Consortium (The 1001 Genomes Consortium 2016). Phenotypic traits used in the present study include flowering time at 10°C (FT10, https://arapheno.1001genomes.org/phenotype/261/), flowering time at 16°C (FT16, https://arapheno.1001genomes.org/phenotype/262/), stomata size (ST, https://arapheno.1001genomes.org/phenotype/750/) and cauline leaf number (CL, https://arapheno.1001genomes.org/phenotype/705/). AraPheno stores 1,163 world-wide *A. thaliana* accessions. We split the 888 European accessions into eight subpopulations of approximately equal sizes (103-119 accessions) (Fig. 1 and Table S1). For ST and CL, the total number of accessions used in our analyses was 240. For both traits, the initial group of 240 accessions was split into two geographic subpopulations, one containing 109 Iberian accessions and the other 131 Swedish accessions. In addition to these traits, we used expression data (Kawakatsu et al. 2016) for 24,175 genes measured in 727 different accessions, available via AraPheno (https://arapheno.1001genomes.org/study/52/). We selected the 665 accessions with full genome sequencing data, and created two subpopulations roughly matching the cauline leaf and stomata size data. The “Swedish” subpopulation contains 70 accessions from Sweden, 2 accessions from Denmark and 2 accessions from Norway, while the second subpopulation, from the Iberian Peninsula contains 83 accessions from Spain and 8 accessions from Portugal. The RNA-seq data have been generated in two distinct batches (Yoav Voichek, personal communication), but accessions from both subpopulations were predominantly present in the second batch, minimizing the risk of batch effects in the analyses.

### 4.2 GWAS

GWAS was performed using a liner mixed model to account for population structure. We used a custom R script (available at https://github.com/arthurkorte/GWAS) implementing a fast approximation of the described in Kang et al. (2010). Significance thresholds were defined using both Bonferroni- and permutation-based thresholds. The Bonferroni threshold was calculated by dividing the significance level (*α* = 0.05) by the number of SNPs with minor allele count greater five in each GWAS run. Permutation-based thresholds were derived from running 100 linear mixed models per phenotype with a random reordering of the phenotypic values (Freudenthal et al. 2019).

### 4.3 Candidate gene enrichment

To look for an enrichment of *a priori* candidate genes, the regions identified as significantly associated with flowering time were been cross-referenced with a list of 306 known flowering time genes (Bouché et al. 2016). All genes within 10 kb of an associated regions were considered. This analysis was conducted with the 74 regions that were associated with flowering time in at least two subpopulations. 22 of these regions overlapped with known flowering time genes. Permutation analysis by re-sampling random regions of the same size across the genome, showed that there is no significant enrichment of candidate genes. Neither changing the window size, nor restricting the analysis to regions that are shared in three or more subpopulation affected this conclusion.

### 4.4 Simulations

In order to simulate data that mimic local and global effects, we use the same subpopulations used for the stomata size and cauline leaf GWAS. We simulated three scenarios:

1. A single marker explaining *x* % of the variance in the full population of 240 accessions;
2. A single marker explaining *x* % of the variance only in the 109 IP accessions, and;
3. A single marker explaining *x* % of the variance only in the 131 Swedish accessions.

In each scenario, the causal marker was chosen randomly from all markers with a minor allele count greater five and set to explain 20%, 15%, 10% and 5% of the phenotypic variance, respectively. To mimic population structure, 1,000 random markers where additionally assigned random small effects that are zero-centered. 1,000 simulated phenotypes were generated for each setting, resulting in a total of 12,000 simulated phenotypes. All simulated data were generated using a custom R script (https://github.com/arthurkorte/GWAS). When the simulated causative marker explained 20% of the phenotypic variation, GWAS performed using all accessions resulted in the detection of this causative marker in 96.4% of the cases, albeit at a high false discovery rate (FDR) of 18.9%. Here, we consider an association as false, if it is more then 100 kb apart from the simulated causal marker. This high FDR dropped dramatically when a more stringent threshold of *p <* 10^*−*9^ or *p <* 10^*−*10^ was applied. Even with this more stringent threshold, a power of 87.6% and 79.4 % was reported, while the FDR dropped to 8.4 % and 4.8 %, respectively. We observed a reduced power in GWAS when using the two different subpopulations (24.6% in IP and and 39% in SW). The reduced detection rate of the marker in IP and SW is caused by a reduced power due to the smaller population size. If the simulations mimic a scenario of a marker having a local effect only, the respective marker was exclusively detected in the respective local subpopulation (42% in SW and 27.4% in IP) and - with a reduced power - in the analyses using all accessions (6.5% and 27%, respectively). Representative GWAS results of the simulated phenotypes are presented in Fig. S10. The analyses of simulations with a reduced effect size of the causative marker led to similar results, albeit at a reduced power (Table S8).

### 4.5 Polygenic overlap

First, we estimated the polygenic overlap among all subpopulations by comparing lists of significant SNPs. Since the comparison of significant SNPs between subsets showed no shared signals, we set a less stringent p-value threshold (*p <* 10^*−*4^) and generated a new list of SNPs for comparing subpopulations. Additionally, we looked at shared significant genomic regions. For this, we summarized all SNPs (*p <* 10^*−*4^) with either r^2^>0.9 or located within a 10 kb window for each subpopulation and compared significant genomic regions. The same procedure has been performed for the respective GWAS results of the subpopulations, as well as with GWAS results from permutations within the respective subpopulation to compare the overlap to the expected overlap in a scenario where no causal markers are present.

Next, we estimated the polygenic overlap using the statistical tool *MiXeR* (Frei et al. 2019), which overcomes the intrinsic problem of detecting the exact location of shared causal variants. In short, a summary table containing SNP information, genomic location, beta estimates, and z-scores for each subpopulation was created and used to estimate the proportion of shared causal SNPs between subsets based on their beta and z-score distributions.

### 4.6 RNA expression data

The available RNA expression data contain transcription values for 24,175 genes. Before performing GWAS on the RNA expression data, we removed TEs and genes that are encoded by the organelle genomes, leaving 23,021 nuclear genes for further analyses. Next, we selected genes where the pseudo-heritability estimate was above 0.5 and a statistical power analysis estimated that the power in GWAS was greater than 0.9 (using the method of Wang and Xu 2019). Heritability was estimated for all genes using the above mentioned implementation of the mixed model. The power of each data set for GWAS was calculated using the *pwr*.*p*.*test* function implemented in the R package *pwr* (R Development Core Team 2008). This filtering led to a set of 2,237 genes for which GWAS was performed in both subpopulations (IP and SW), as well as in the combined population (ALL). We only considered markers with a minor allele count of more than five in the respective subpopulation. Given the amount of tests we performed, we used a very stringent multiple-testing threshold of *p <* 10^*−*10^ to term an association as significant, but similar results have been reproduced with threshold ranging from *p <* 10^*−*8^ to *p <* 10^*−*12^. Significant associations were grouped into regions, if they occur within 50 kb of each other. A summary of the number of associated markers and regions for all analysed genes as well as summary statistics are attached in File S2. Genes showing inflated GWAS results (which quite often co-occurs with a non-normal distribution of the expression values), have been filtered out, if the number of associated genomic region was greater then three in either the IP or SW subpopulation. This procedure left us with a set of 1,982 genes. GWAS results from these selected genes were analysed in more detail and the complete workflow of the analysis is displayed in Fig. S11. We identified 227 genes displaying an association in both the IP and SW subpopulation. For 135 of them, the same genomic region was associated in both subpopulations, while for 92 genes different genomic regions have been associated in the two subpopulations (File S3). To prevent genes from being assigned as locally regulated in both subpopulations, those genes have not been considered as genes displaying a local regulation. Still, these genes show the same pattern of *cis*-*vs. trans*-regulation observed for genes with a specific local association. Genes, where the same genomic region was associated, were defined as genes having a global genetic regulation, if the same significant marker in both subpopulations was associated (110, File S4), while genes where the same region but different markers are associated in the subpopulations (25, File S5), were classified as genes showing potential allelic heterogeneity in their regulation. Next, genes that show an association only in one and not the other subpopulation were defined as genes that are under distinct local regulation. This led to the identification of 377 genes displaying an association only in IP and 176 genes displaying an association only in SW. Now, we filtered for genes, where the respective p-value was lower in the analysis of the respective subpopulation compared to the results of the combined population, as we argue that a true local association should be more significantly associated in the respective local subpopulation. Additionally, we also excluded genes, where different regions have been associated in the analysis of the combined population compared to the analysis of respective local subpopulation, to generate a high confidence list of genes with a distinct regulation in only one subpopulation. This procedure led to a set of 118 genes displaying a specific local association only in the Iberian subpopulation (File S6) and 64 genes displaying a specific local association only for the Scandinavian subpopulation (File S7). For significant associations in these three groups of genes, having the same association in both subpopulations, a specific local association only in IP or a specific local association only in SW, we verified, if the respective associated SNPs were in *cis*, aka the same genomic region where the gene is located, or in *trans*. Here, we defined a *cis*-association, by a maximum distance of the associated markers to the respective gene of 100 kb. As a control, we also performed the same analysis described above with two random, non-local population of 91 and 74 accessions, respectively. These random populatilons have been sampled from the merged population of 165 accessions. The respective workflow and numbers are presented in Fig. S15. Note, that we started out with the same set of 2,237 genes used previously, but here the removal of genes showing inflated results, led to a set of 2,087 genes included in the analysis.

## 5 Acknowledgements

We thank Yoav Voichek for providing data for the batch effects of the RNAseq data. Additional thanks to Pieter Clauw and Pamela Korte for detailed comments on the manuscript.

## 6 Data Accessibility

All data and scripts used are publicly available and the respective locations and links are provided in Methods section.

## 7 Author Contributions

MN and AK planned and designed the study. WLA, SR and AK performed the statistical analyses. All authors wrote, read and approved the manuscript.

## 8 Supplementary Figures

**Supplementary Fig 1:**
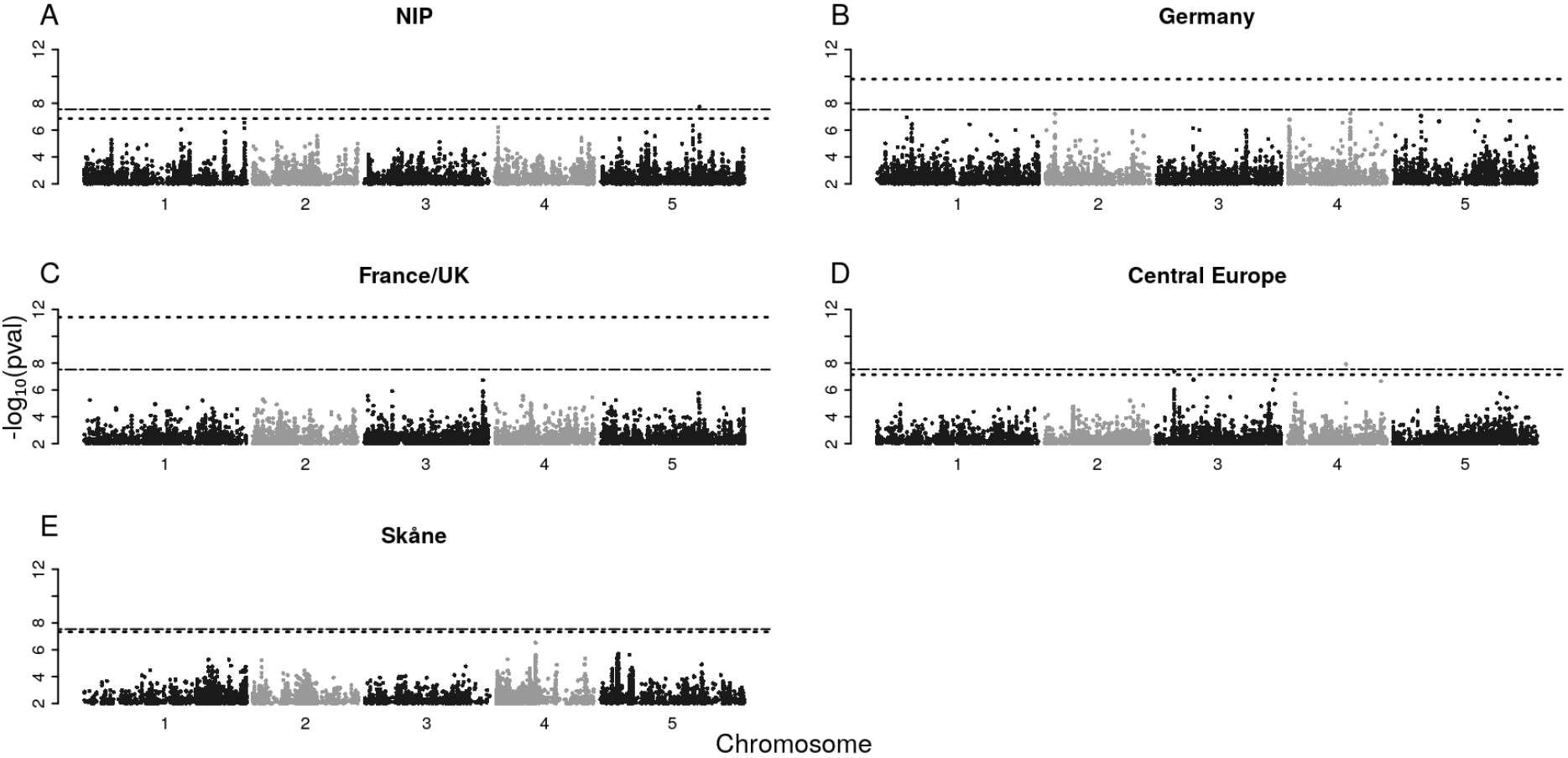
Manhattan plot for the GWAS results of flowering time in five different subpopulations. Dashed lines and dash-dotted lines indicate 5% permutation-based and Bonferroni threshold, respectively.

**Supplementary Fig 2:**
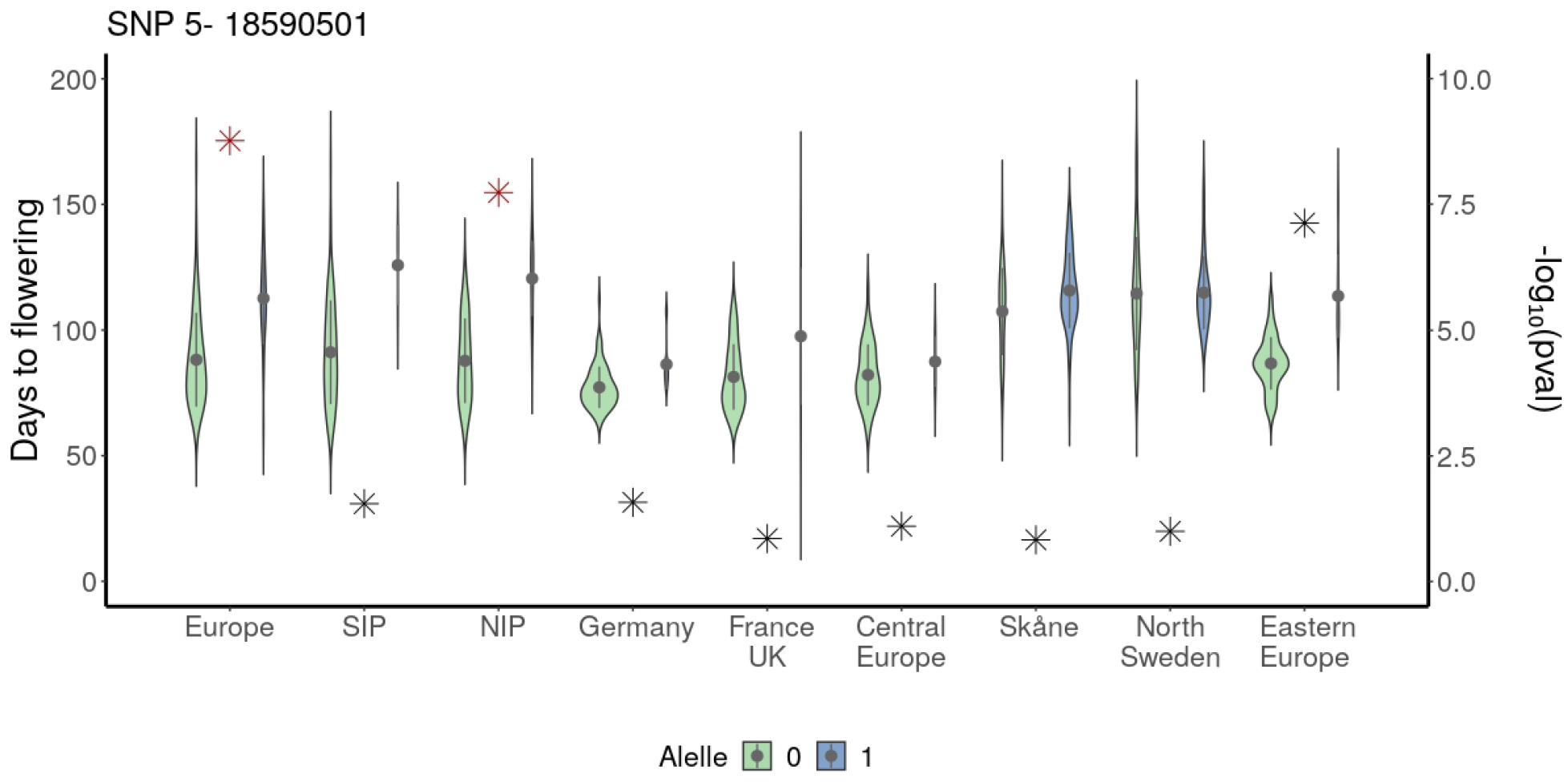
Violin plots comparing flowering time between accessions carrying reference and alternative allele for the SNP 5:18590501. Stars represent the -log_10_ of the p-value and a red colored star indicates a significant association.

**Supplementary Fig 3:**
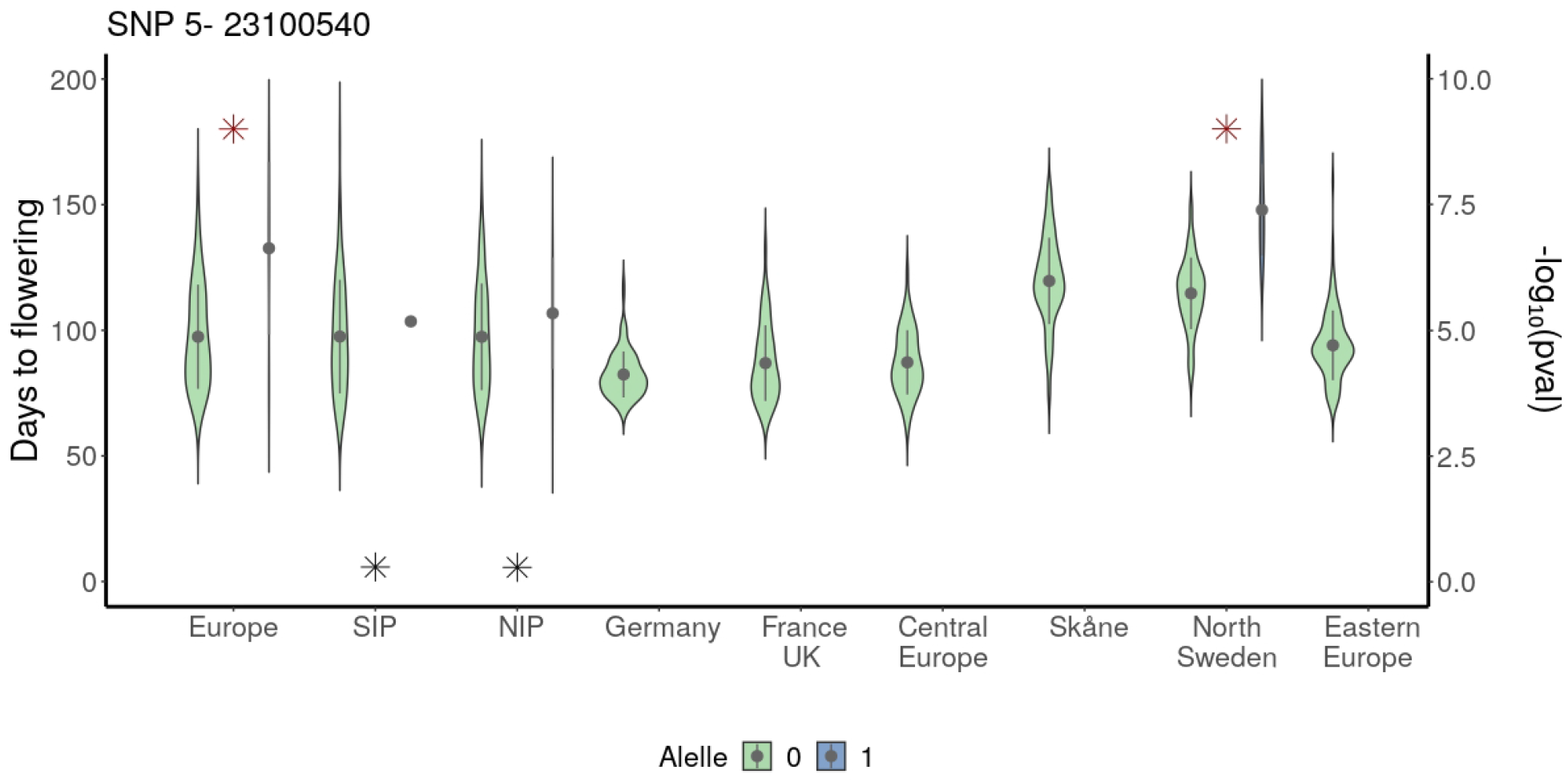
Violin plots comparing flowering time between accessions carrying the reference or alternative allele for SNP 5:23100540. Stars represent the -log_10_ of the p-value and a red colored star indicates a significant association.

**Supplementary Fig 4:**
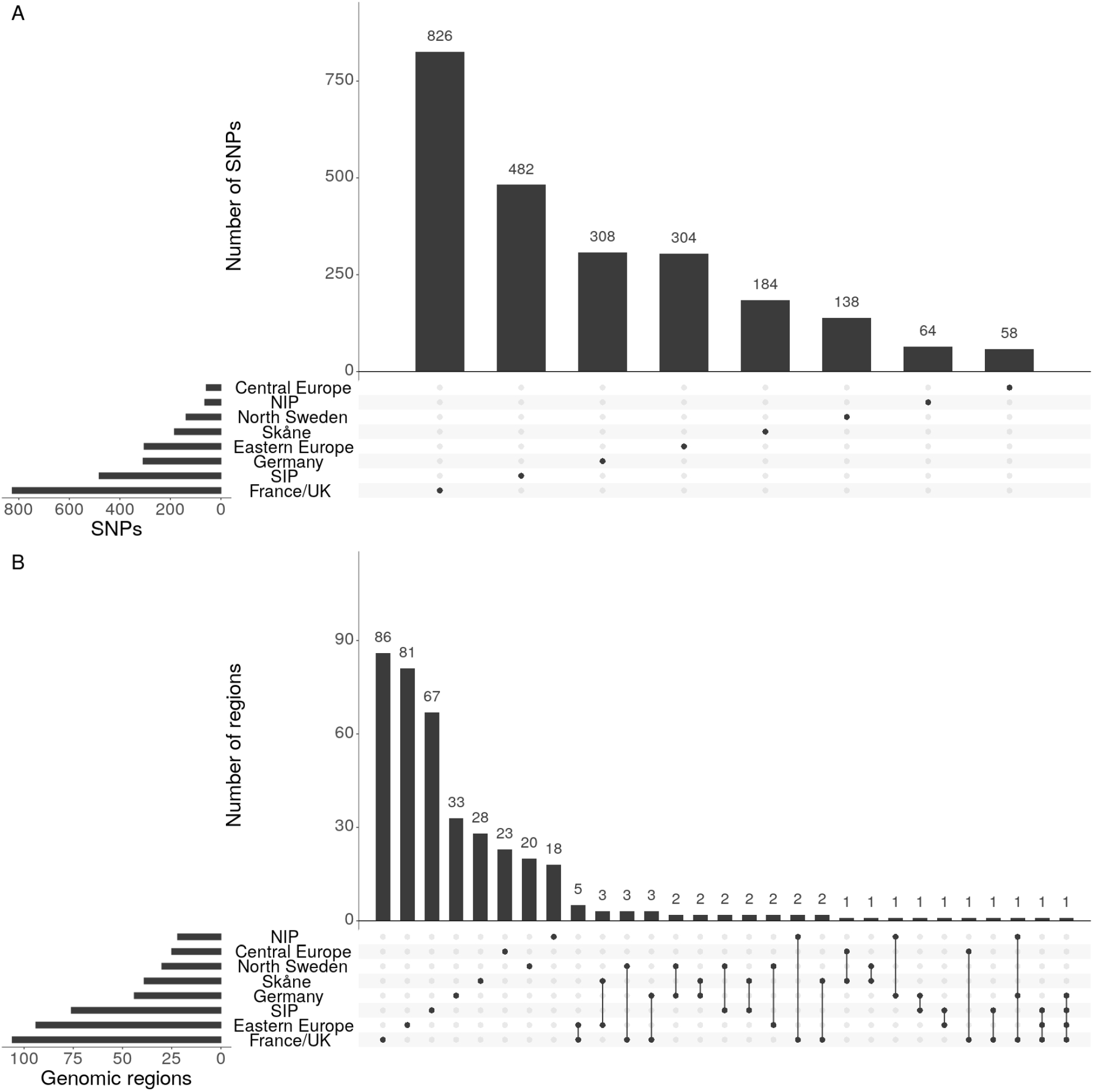
Sharing of sub-significant (*p <* 10^*−*4^) associations for permuted phenotypes. (**A**) Histogram of the number of associated SNPs in each subpopulation and shared between pairs of subpopulations. (**B**) Histogram of the number of associated regions in each subpopulation and shared between pairs of subpopulations.

**Supplementary Fig 5:**
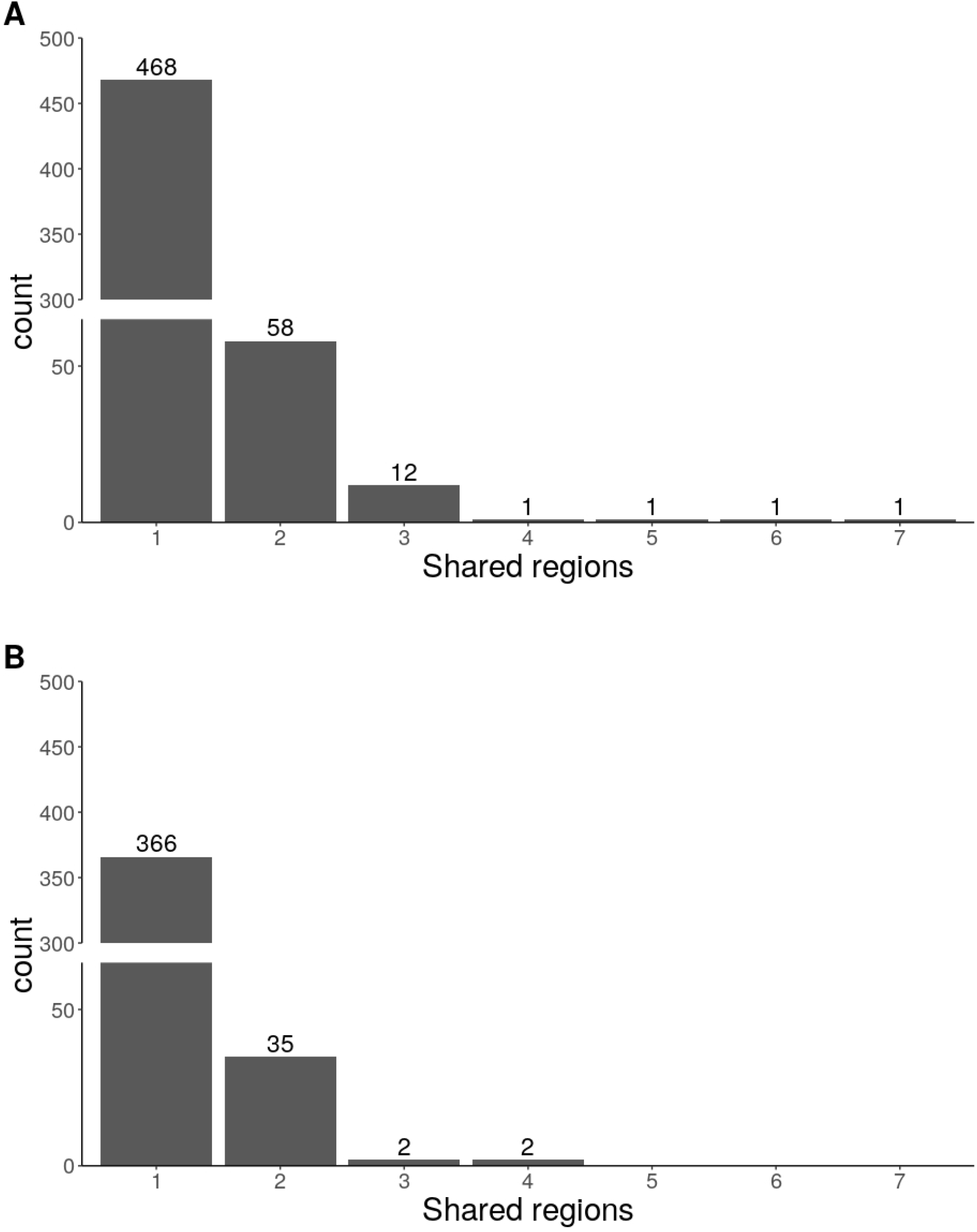
Bar plot showing the amount of regions that are associated across multiple sub-populations. (**A**) Analyses of FT10. (**B**) Analyses of permuted phenotypes.

**Supplementary Fig 6:**
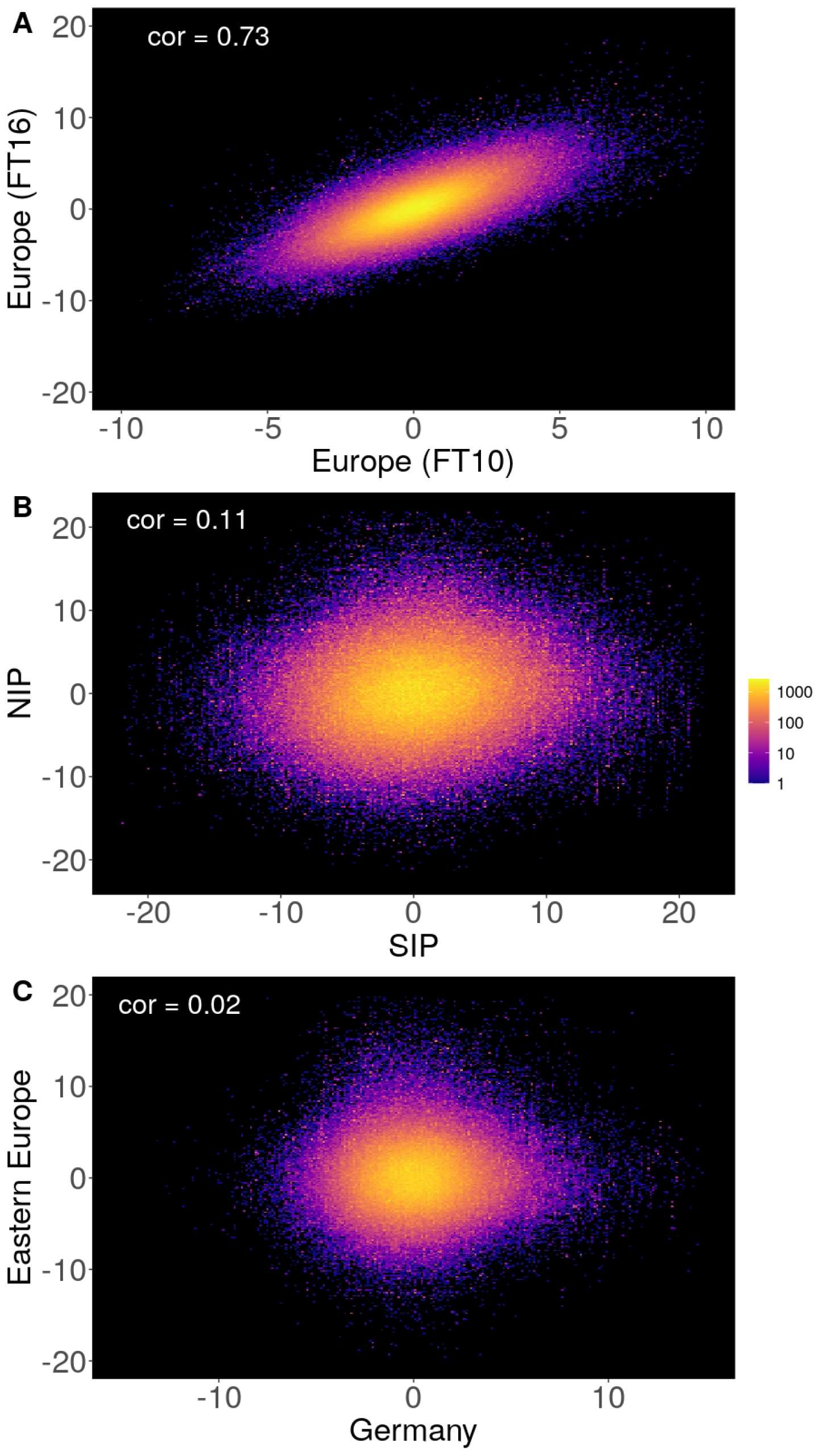
Density plots comparing effect sizes estimates using *MiXeR*. (**A**) The comparison between flowering time at 10°C and 16 °C in the complete European population. (**B**) Comparison of FT10 between NIP and SIP subpopulations. (**C**) Comparison of FT10 between the Eastern Europe and German subpopulations.

**Supplementary Fig 7:**
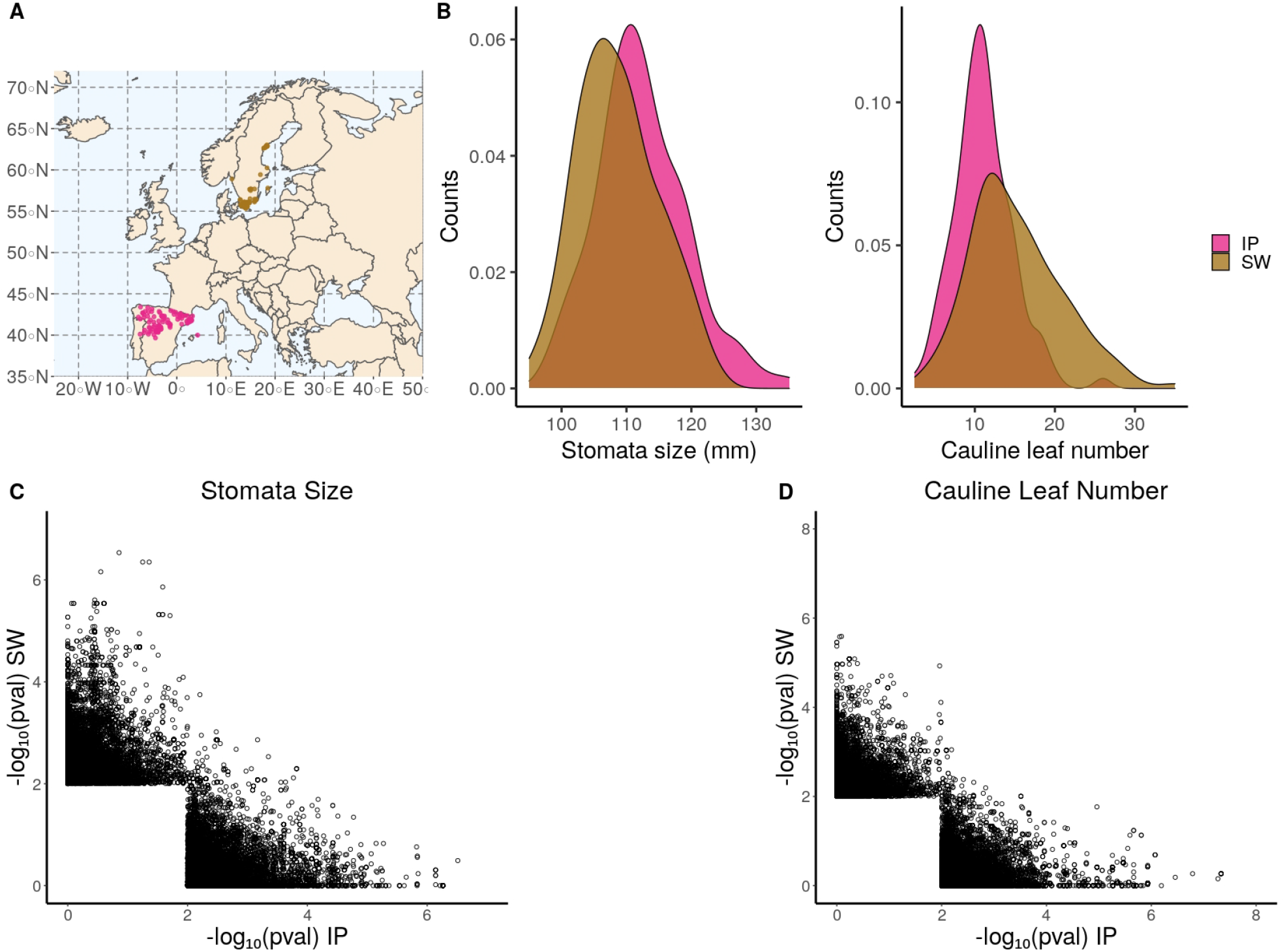
Analyses of stomata size and cauline leaf number.(**A**) Geographic distribution of the used 240 *A. thaliana* accessions.(**B**) Phenotypic distribution of stomata size and cauline leaf number in the Iberian (IP) and Swedish (SW) subpopulation.(**C-D**) Correlation plots of the -log_10_ p-values from the GWAS results obtain from the analyses within the different subpopulations for ST and CL. Marker with a p-value > 0.01 where removed for plotting from the respective subpopulation.

**Supplementary Fig 8:**
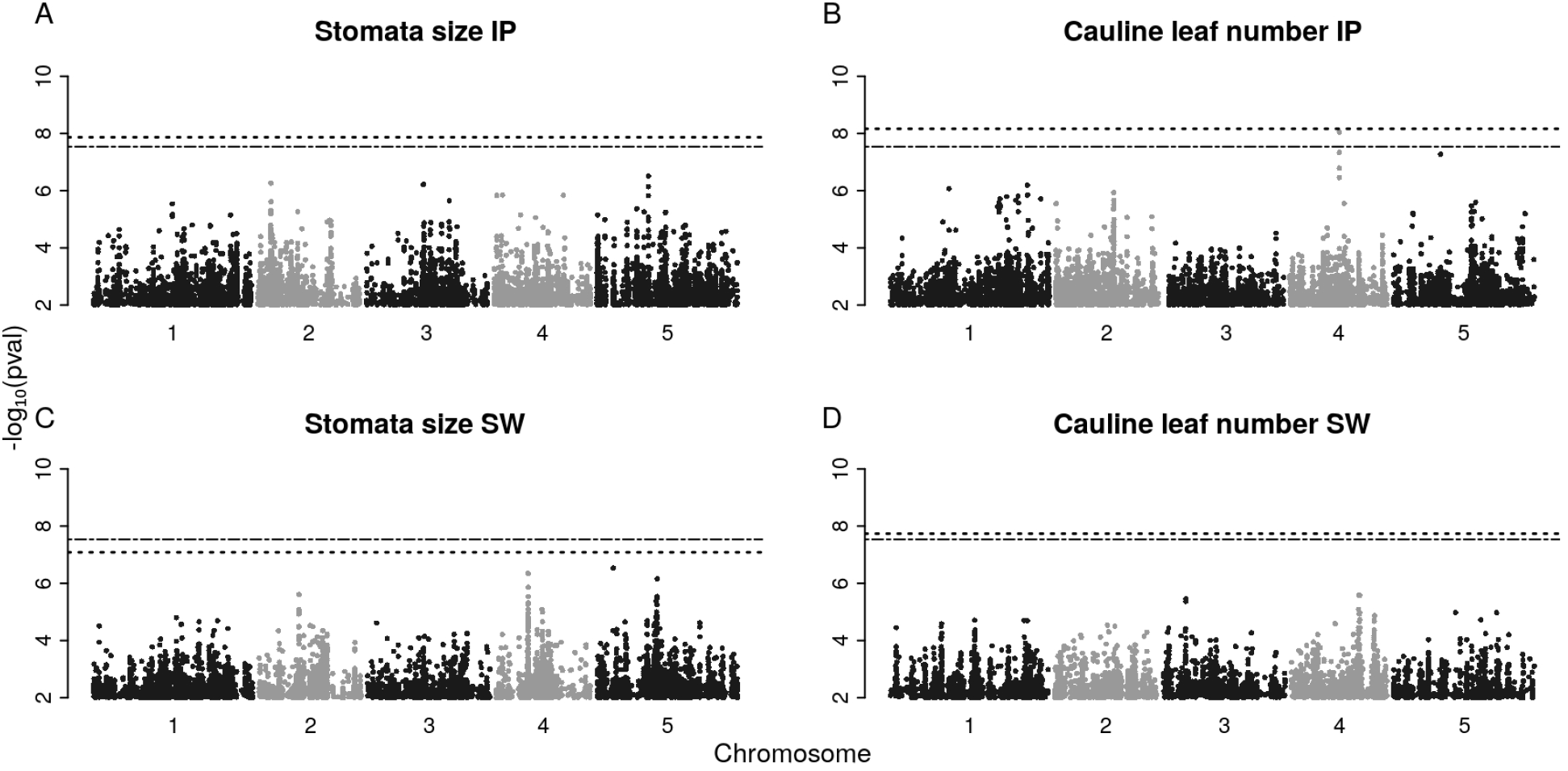
Manhattan plots of GWAS results from the analyses of ST and CL in the SW and IP subpopulations.

**Supplementary Fig 9:**
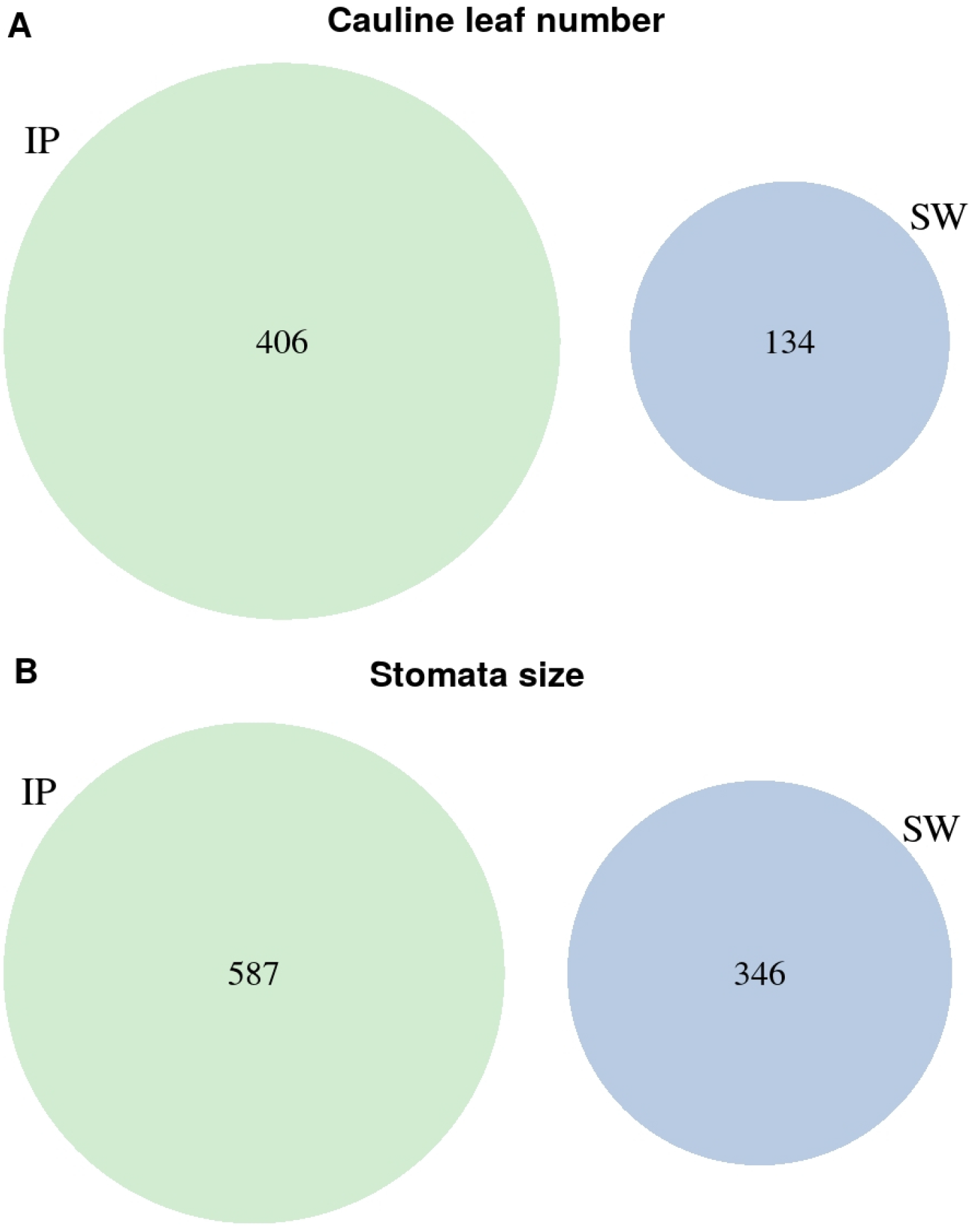
Venn diagram of associated markers (*p <* 10^*−*4^) between the IP and SW subpopulations for ST (A) and CL (B).

**Supplementary Fig 10:**
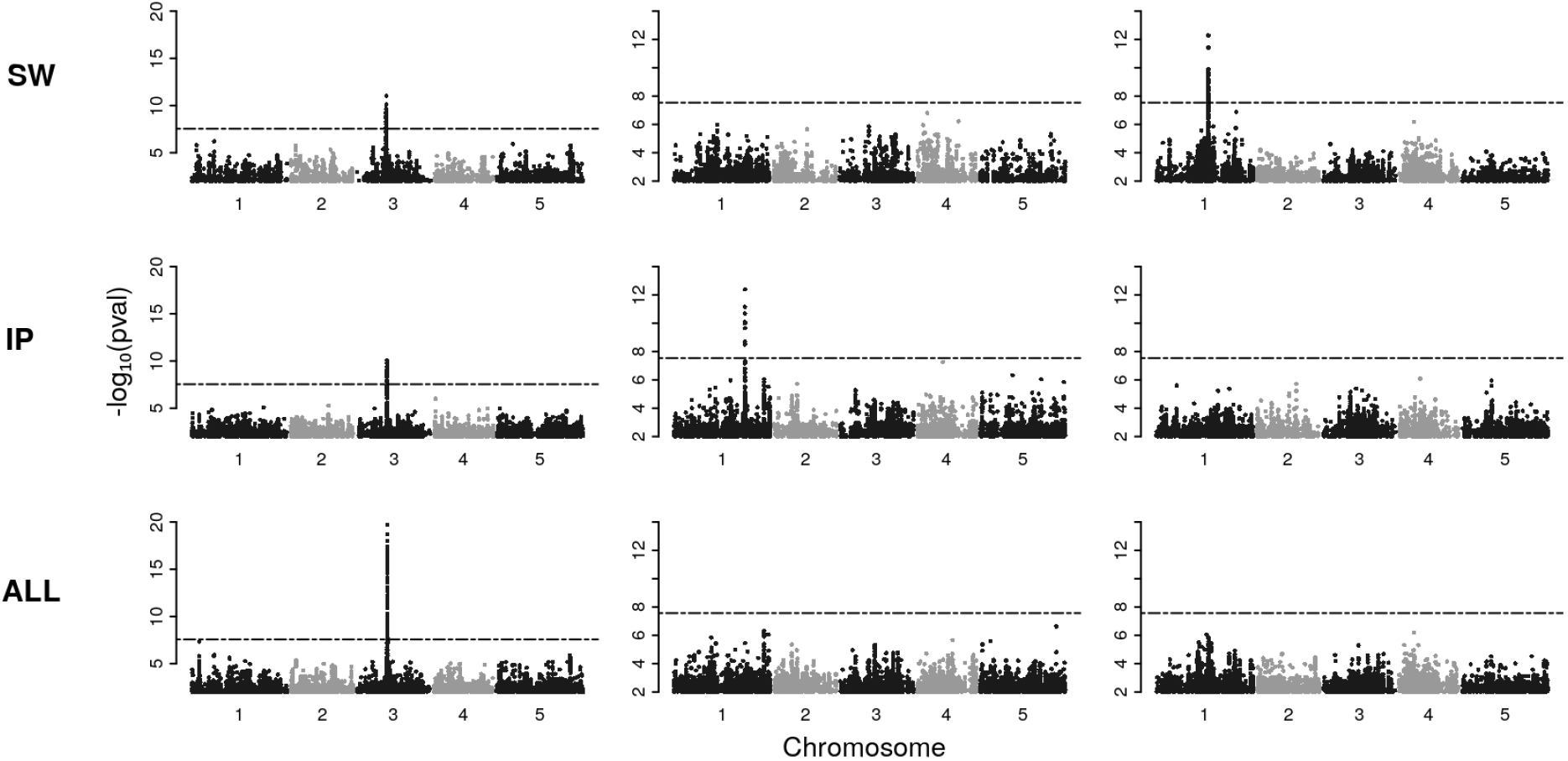
Manhattan plots of GWAS results from three different simulations. The causative markers were simulated to have an effect in all accessions (left panel), only in IP (middle panel) or only in SW (right panel). The respective population for GWAS are displayed in the different rows, where for the results in the top row, the SW subpopulation has been used, the IP subpopulation has been used to generate the results in the middle row and the bottom row displays the results in the merged population of 240 accessions. Dashed lines indicate the Bonferroni threshold used in the simulations.

**Supplementary Fig 11:**
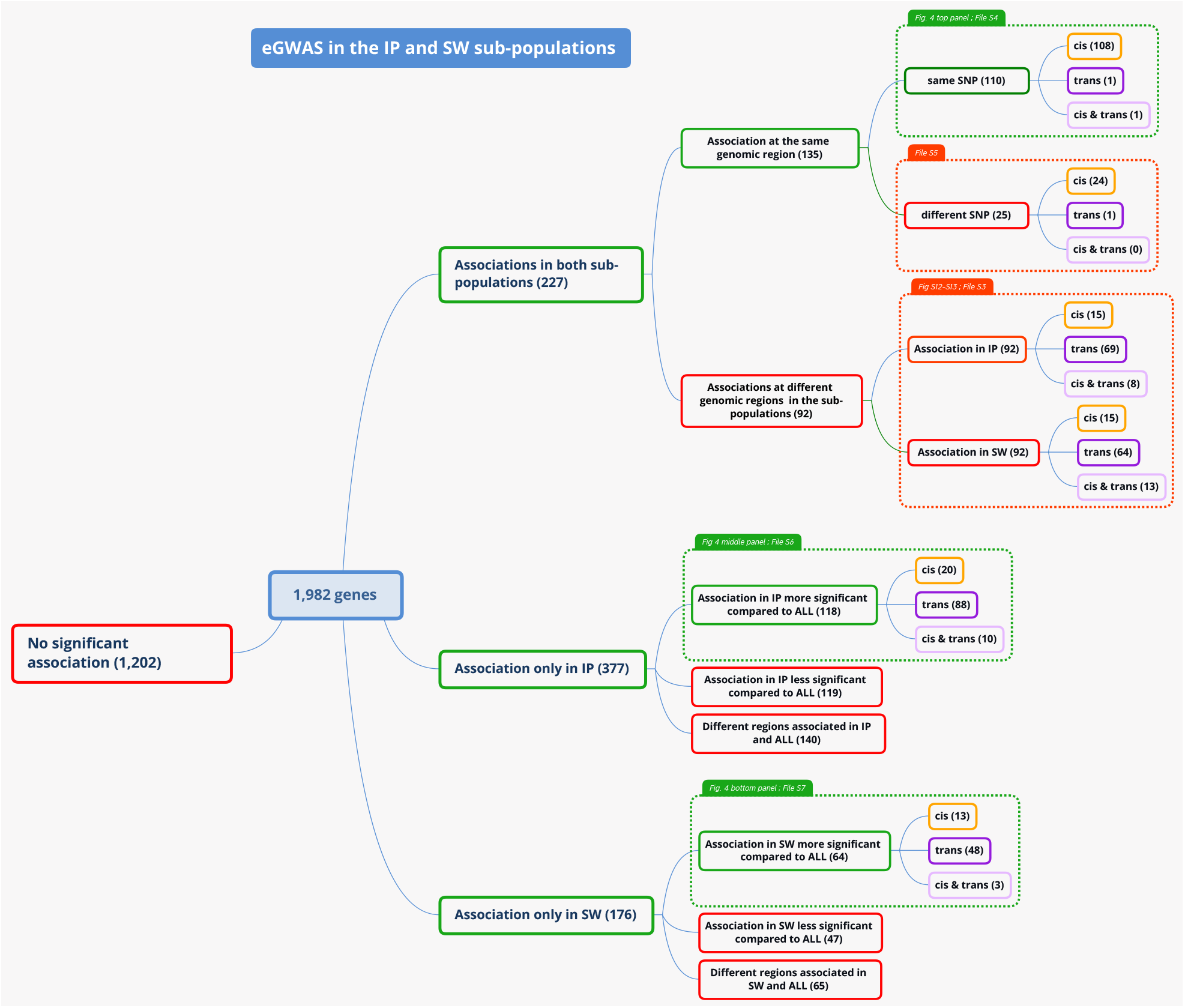
Graphical representation of the workflow and the different classes of associations found in the eGWAS for the IP and SW subpopulations at a threshold of *p <* 10^*−*10^.

**Supplementary Fig 12:**
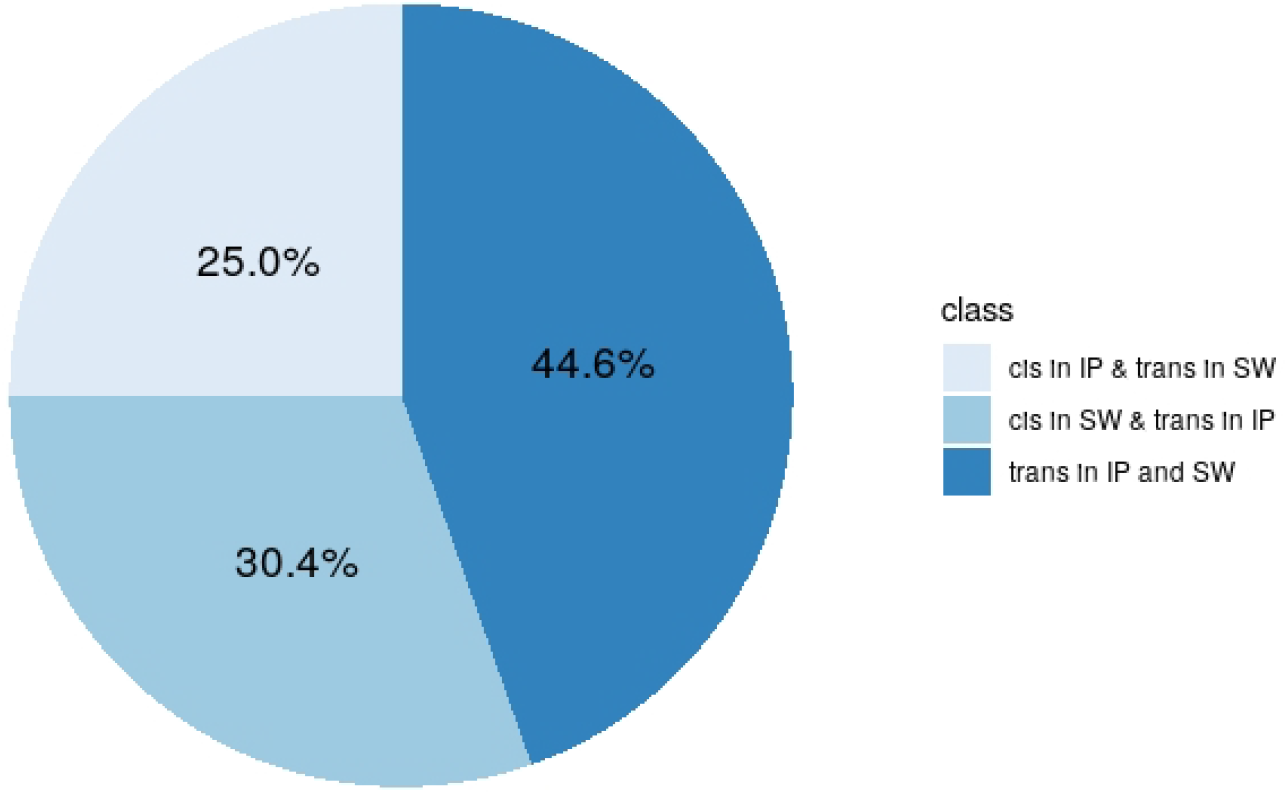
Summarized GWAS results for the analyses of RNA expression data in *A. thaliana* for genes that have associations at different regions in IP and SW. The pie chart displays the amount of genes having a *cis*-association in IP and a *trans*-association in SW, a *cis*-association in SW and a *trans*-association in IP or different *trans*-associations in both subpopulations.

**Supplementary Fig 13:**
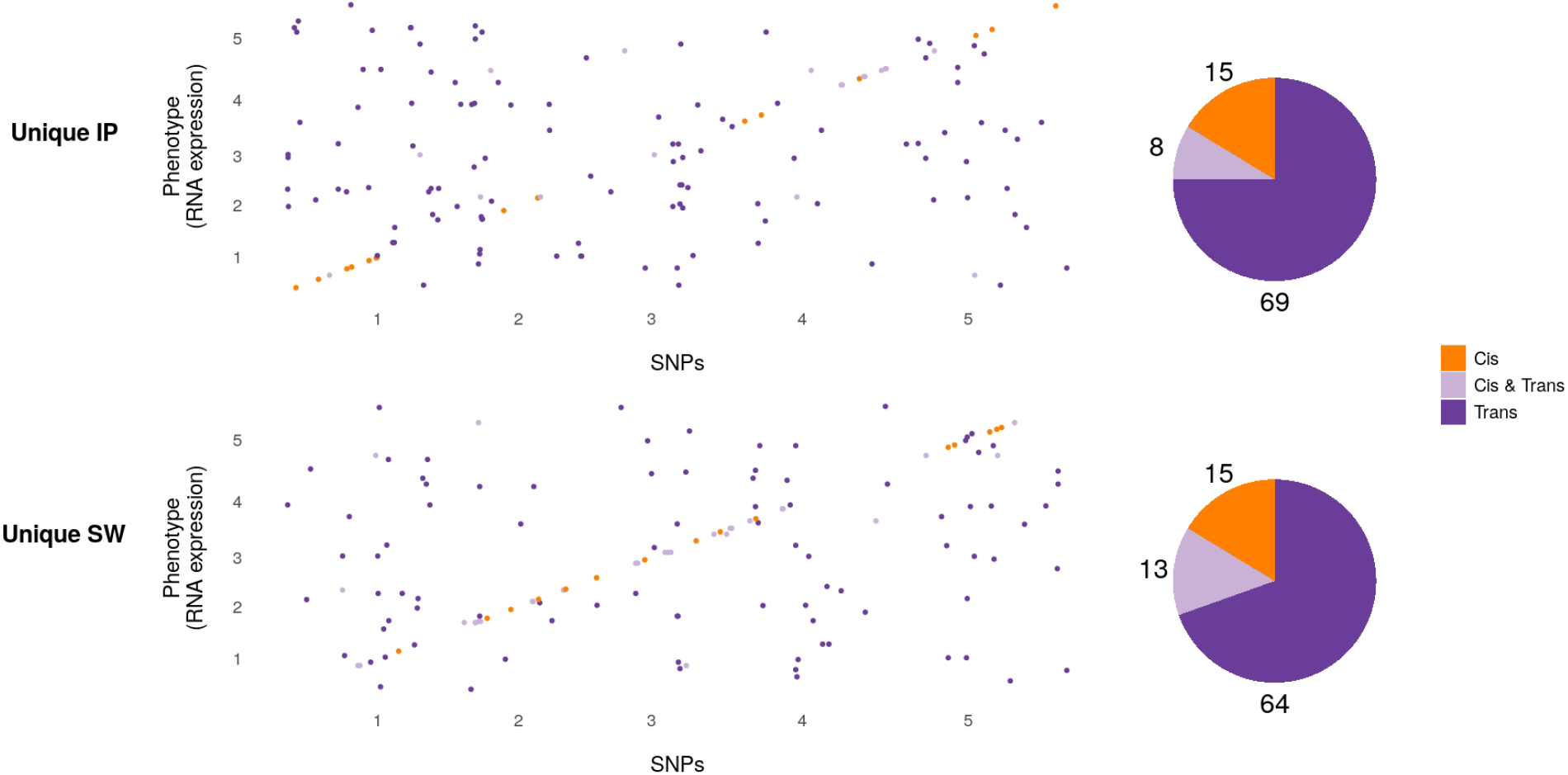
Summarized GWAS results for the analyses of RNA expression data in *A. thaliana* for genes that have associations at different regions in IP and SW. Scatter plots show the genomic location of the respective associated markers per gene. Top panel shows the amount of *cis* and *trans* associations in IP, and the bottom panel shows the location of the associations for the same genes in SW. *Cis*-regulatory variants are colored in orange, while variants in *trans* are shown in purple. Pie charts display the amount of genes per class that have *cis, cis* and *trans* or only *trans* associations.

**Supplementary Fig 14:**
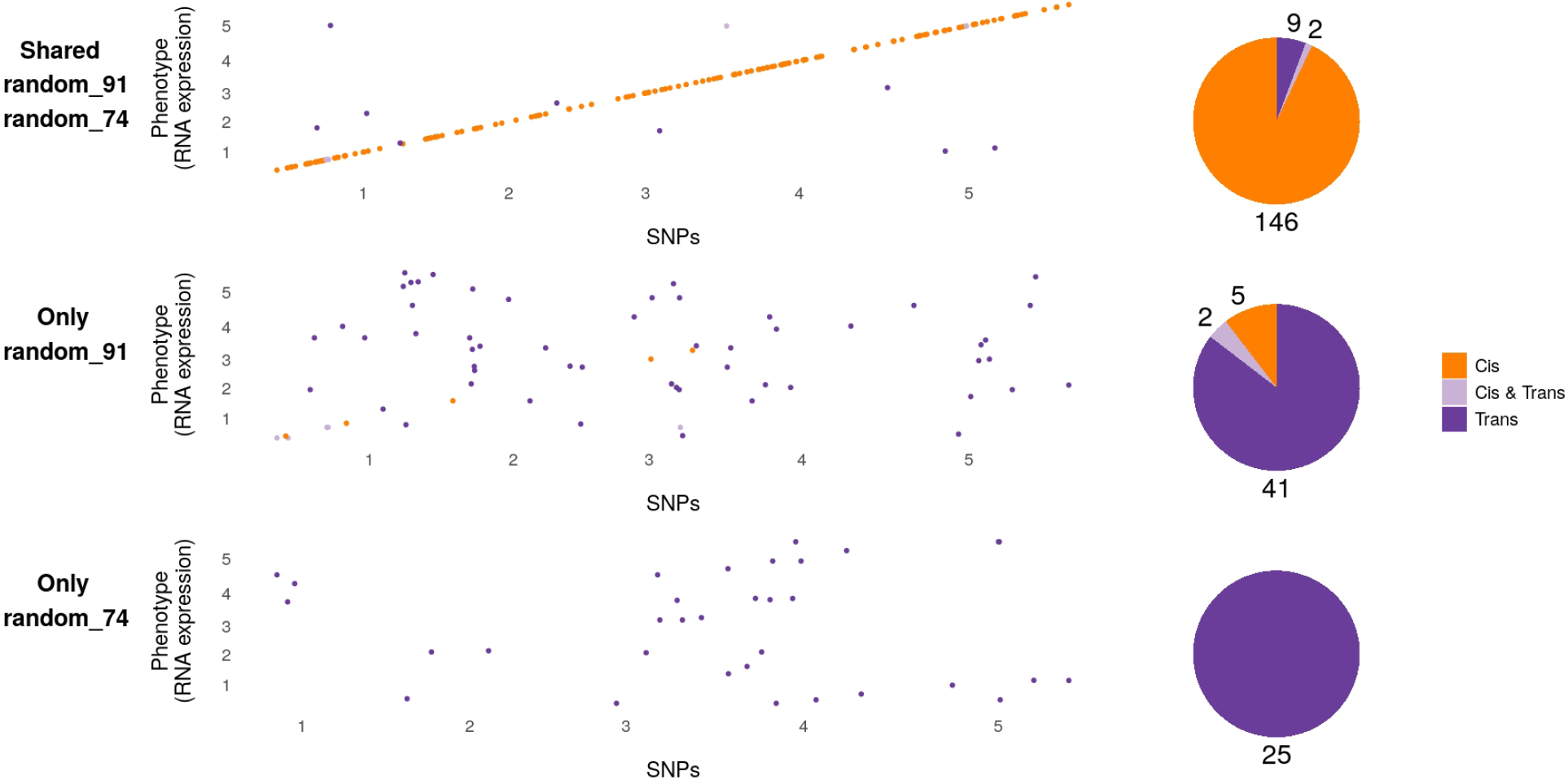
Summarized GWAS results for the analyses of RNA expression data in *A. thaliana* in random subpopulations. Genes are grouped in three categories: 1) Shared random_91/random_74, where the same association for a gene is recapitulated in the GWAS of both subpopulations. 2) Only random_91, where a significant association is only found using the random subpopulation containing 91 accessions. 3) Only random_74, where a significant association is only found using the random subpopulation containing 74 accessions. Scatter plots show the genomic location of the respective associated markers per gene for each class, where *cis*-regulatory variants are colored in orange, while variants in *trans* are shown in purple. Pie charts display the amount of genes per class that have *cis, cis* and *trans* or only *trans*-associations.

**Supplementary Fig 15:**
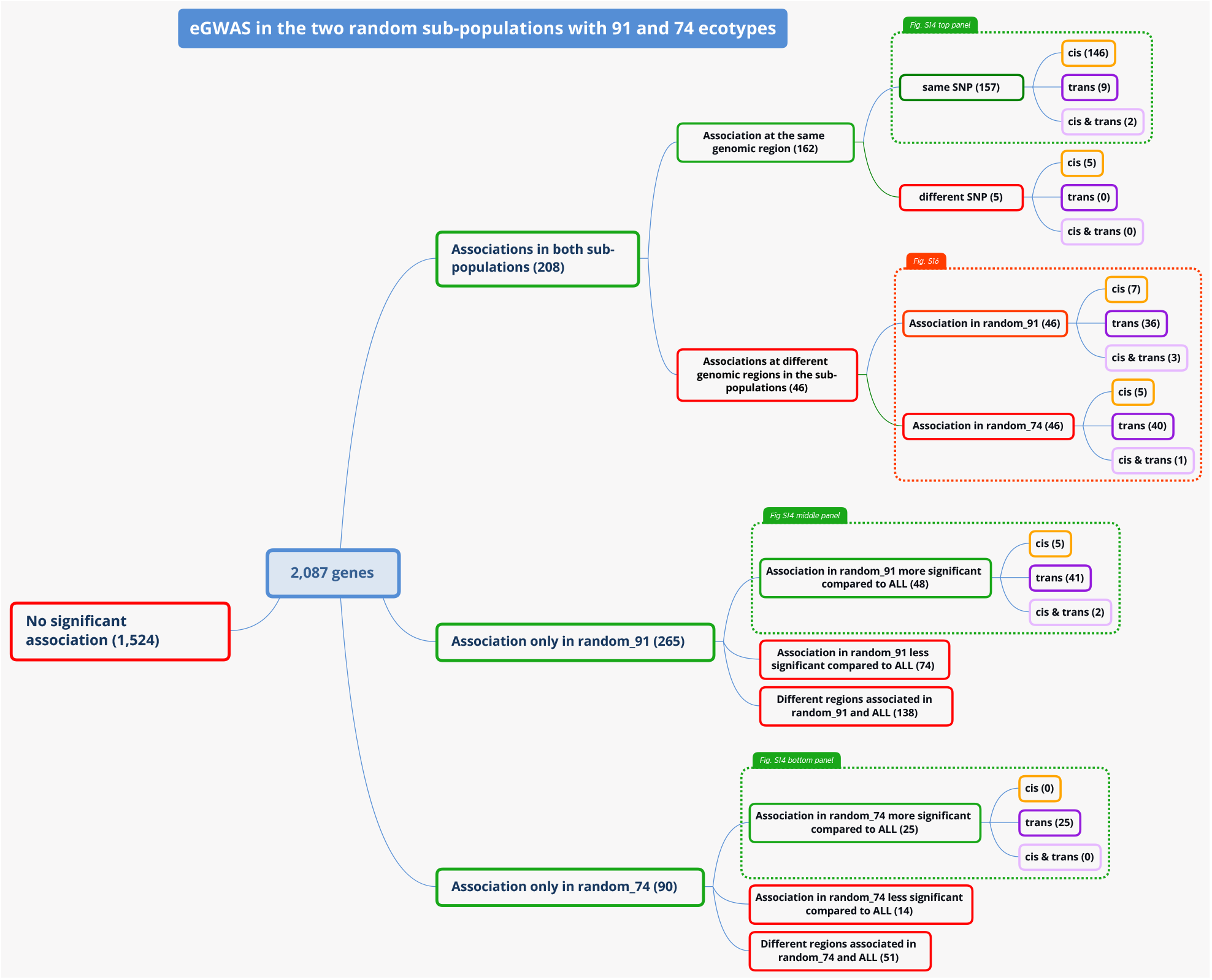
Graphical representation of of the different classes of associations found in the eGWAS for two random subpopulations of 91 and 74 accessions at a threshold of *p <* 10^*−*10^.

**Supplementary Fig 16:**
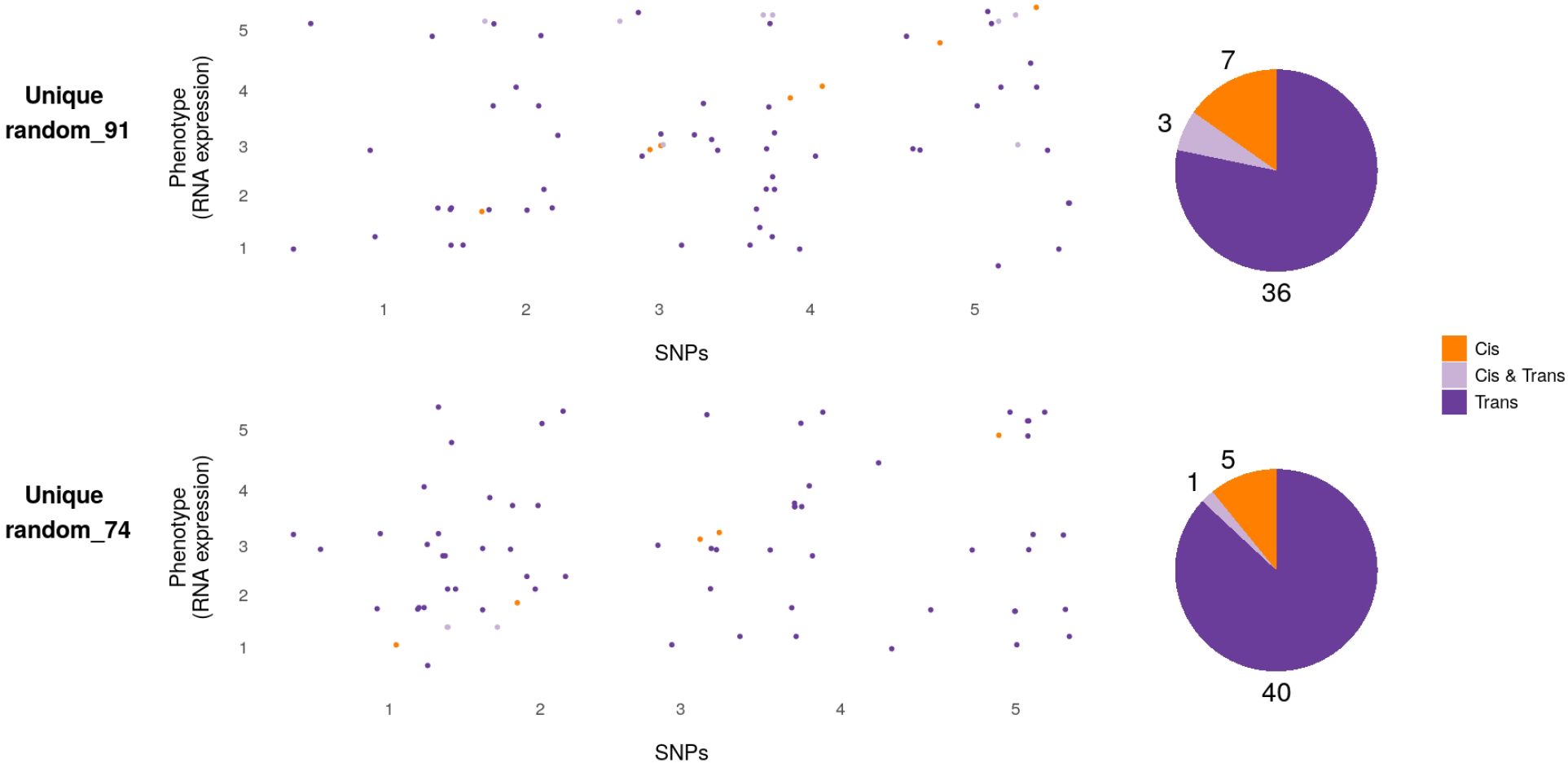
Summarized GWAS results for the analyses of RNA expression data in *A. thaliana* in random subpopulations for genes that have associations at different regions in both random subpopulations. Scatter plots show the genomic location of the respective associated markers per gene. Top panel shows the amount of *cis* and *trans*-association in random_91, and the bottom panel show the location of the associations for the same genes in random_74. *Cis*-regulatory variants are colored in orange, while variants in *trans* are shown in purple. Pie charts display the amount of genes per class that have *cis, cis* and *trans* or only *trans*-associations.

## 9 Supplementary Tables

**Supplementary Table 1:** Geographic location of the European subpopulations.

| Subsets | No. accessions | min_lat | max_lat | min_lon | max_lon | $\widehat{h}_2^a$ | power |
| --- | --- | --- | --- | --- | --- | --- | --- |
| SIP | 107 | 36.52 | 41.48 | -8.54 | 4.25 | 0.99 | 1.00 |
| NIP | 108 | 41.50 | 47.45 | -7.80 | 6.13 | 0.91 | 0.99 |
| Germany | 107 | 48.39 | 55.67 | 8.00 | 13.73 | 0.99 | 1.00 |
| France/UK | 107 | 47.50 | 57.97 | -5.98 | 7.50 | 0.95 | 0.99 |
| Central Europe | 106 | 37.30 | 49.37 | 6.08 | 17.31 | 0.66 | 0.99 |
| Skåne | 119 | 55.38 | 56.10 | 13.10 | 14.78 | 0.91 | 0.99 |
| Northern Sweden | 118 | 56.10 | 68.80 | 6.19 | 18.52 | 0.75 | 0.99 |
| Eastern Europe | 116 | 37.07 | 61.36 | 38.28 | 38.28 | 0.91 | 0.99 |
| Europe | 888 | 36.52 | 68.80 | -8.54 | 38.28 | 0.86 | 1.00 |

<sup>a</sup> $\widehat{h}_2$ : pseudo-heritability estimate

**Supplementary Table 2:** Number of significant SNPs after Bonferroni and permutation-based threshold in the different subpopulations

| Subsets | Bonferroni_Threshold | Sig_Bonferroni | Permutation_Threshold | Sig_Permutation |
| --- | --- | --- | --- | --- |
| Europe | 1.1e-08 | 29 | 1.7e-09 | 2 |
| SIP | 2.5e-08 | 33 | 2.6e-09 | 2 |
| NIP | 2.6e-08 | 4 | 1.4e-07 | 4 |
| Germany | 2.7e-08 | 14 | 1.6e-10 | 0 |
| France and UK | 2.8e-08 | 0 | 3.8e-12 | 0 |
| Central Europe | 2.6e-08 | 1 | 7.3e-08 | 12 |
| Skåne | 2.7e-08 | 0 | 4.6e-08 | 0 |
| Northern Sweden | 2.7e-08 | 26 | 1.2e-08 | 2 |
| Eastern Europe | 2.6e-08 | 1 | 1.3e-10 | 0 |

**Supplementary Table 3:** P-value and minor allele frequency of the SNPs that are significant in the European set (after Permutation-based threshold) in the different subpopulations

| subpopulation | 1:24339560<sup>a</sup> | 10 kb window<sup>b</sup> |  | 5:18590501<sup>a</sup> | 10 kb window<sup>b</sup> |  |
| --- | --- | --- | --- | --- | --- | --- |
|  |  | SNP | Pval (MAF) |  | SNP | Pval (MAF) |
| Europa | **2.39e-10** (0.44) |  |  | **1.71e-09** (0.20) |  |  |
| SIP | 1.68e-02 (0.45) | 1:24339614 | 9.88e-04 (0.30) | 2.93e-02 (0.28) | 5:18605212 | 3.54e-03 (0.47) |
| NIP | 1.25e-02 (0.35) | 1:24338003 | 1.40e-04 (0.30) | **1.83e-08** (0.14) | 5:18590501 | **1.83e-08** (0.14) |
| Germany | 3.08e-02 (0.28) | 1:24346295 | 2.82e-04 (0.07) | 2.58e-02 (0.06) | 5:18565932 | 1.49e-03 (0.17) |
| France/UK | 6.99e-04 (0.41) | 1:24342759 | 4.41e-05 (0.40) | 1.43e-01 (0.05) | 5:18606201 | 1.00e-03 (0.4) |
| Central Europe | 7.40e-02 (0.46) | 1:24329854 | 6.70e-03 (0.05) | 8.15e-02 (0.05) | 5:18590327 | 5.20e-03 (0.34) |
| Skåne | 5.1e-02 (0.24) | 1:24320374 | 9.34e-03 (0.27) | 1.51e-01 (0.33) | 5:18585844 | 2.38e-03 (0.07) |
| Northern Sweden | 3.11e-01 (0.20) | 1:24348634 | 1.89e-02 (0.15) | 1.03e-01 (0.48) | 5:18590741 | 1.19e-04 (0.14) |
| Eastern Europe | 2.61e-01 (0.44) | 1:24325815 | 3.47e-03 (0.17) | 7.43e-08 (0.08) | 5:18590501 | 7.43e-08 (0.08) |

<sup>a</sup> "chromosome:position"
<sup>b</sup> the most significant marker in a 10 kb window around the lead SNP is reported for each subpopulation

**Supplementary Table 4:** Shared SNPs between subpopulations at significance level of *p <* 10^*−*4^

| Associated marker<sup>a</sup> | 1:29186215 | 1:29199833 | 3:20379636 | 4:6781375 | 5:18589998 | 5:18590247 | 5:18590501 | 5:18590591 |
| --- | --- | --- | --- | --- | --- | --- | --- | --- |
| Candidate gene<sup>b</sup> |  |  | SMZ<sup>c</sup> |  | DOG1<sup>d</sup> | DOG1<sup>d</sup> | DOG1<sup>d</sup> | DOG1<sup>d</sup> |
| SIP | 3.75e-05 (0.06) | 3.75e-05 (0.06) | 6.22e-01 (0.04) | 1.15e-02 (0.02) | 1.38e-01 (0.04) | 2.92e-02 (0.03) | 2.92e-02 (0.03) | 2.92e-02 (0.03) |
| NIP | 4.80e-01 (0.12) | 4.80e-01 (0.12) | 2.51e-01 (0.06) | 9.45e-01 (0.06) | 1.83e-08 (0.14) | 1.83e-08 (0.14) | 1.83e-08 (0.14) | 1.83e-08 (0.14) |
| Germany | 1.55e-01 (0.10) | 1.04e-01 (0.10) | 9.06e-05 (0.07) | 3.88e-01 (0.07) | 8.28e-03 (0.05) | 2.73e-01 (0.05) | 2.58e-02 (0.05) | 6.30e-01 (0.03) |
| France/UK | 7.34e-01 (0.29) | 7.34e-01 (0.29) | 4.52e-01 (0.20) | 7.75e-05 (0.08) | 1.74e-03 (0.019) | 1.86e-02 (0.03) | 1.43e-01 (0.05) | 5.53e-03 (0.03) |
| Central Europe | 5.90e-01 (0.03) | 5.9e-01 (0.03) |  | 2.70e-02 (0.13) | 2.60e-01 (0.05) | 4.60e-01 (0.05) | 8.15e-02 (0.05) |  |
| Skåne | 8.55e-01 (0.25) | 8.55e-01 (0.25) | 4.35e-02 (0.10) | 4.44e-01 (0.45) | 8.00e-01 (0.23) | 9.86e-02 (0.32) | 1.51e-01 (0.33) | 1.80e-01 (0.37) |
| Northern Sweden | 4.61e-01 (0.5) | 4.78e-01 (0.5) | 4.49e-01 (0.43) | 7.04e-01 (0.36) | 5.12e-01 (0.34) | 4.59e-01 (0.48) | 1.03e-01 (0.48) | 5.75e-10 (0.36) |
| Eastern Europe | 2.85e-05 (0.05) | 2.85e-05 (0.05) | 9.29e-05 (0.05) | 9.62e-05 (0.06) | 7.43e-08 (0.08) | 1.05e-06 (0.09) | 7.43e-08 (0.08) | 7.43e-08 (0.08) |

<sup>a</sup> represented as "chromosome:position"<sup>b</sup> known flowering time gene within a 10 kb window around the associated marker using a list of 306 flowering time genes from Bouché et al. 2016<sup>c</sup> SMZ (*SCHLAFMÜTZE*), Mathieu et al. 2009<sup>d</sup> *DOG1* (*DELAY OF GERMINATION*), Huo et al. 2016

**Supplementary Table 5:** Overlap of candidate genes with shared genomic regions

| Region<sup>a</sup> | 1:(26151612–26570642) | 4:(192421–572878) | 5:(22786643–23605491) |
| --- | --- | --- | --- |
| Candidate gene<sup>b</sup> | *CDF5*<sup>c</sup> | *FRI*<sup>d</sup>*LIF2*<sup>e</sup>*MED12*<sup>f</sup> | *COL5*<sup>g</sup>*MSI*<sup>h</sup>*VIN3*<sup>i</sup>*ZTL*<sup>j</sup> |
| SIP | 0 | 0 | 1 |
| NIP | 1 | 1 | 1 |
| Germany | 1 | 1 | 1 |
| France/UK | 1 | 1 | 1 |
| Central Europe | 1 | 0 | 1 |
| Skåne | 0 | 1 | 0 |
| North Sweden | 1 | 1 | 1 |
| East Europe | 1 | 0 | 1 |

<sup>a</sup> represented as "chromosome:(start–stop)"<sup>b</sup> known flowering time gene within the detected region using a list of 306 flowering time genes from Bouché et al. 2016<sup>c</sup> *CDF5* (*CYCLING DOF FACTOR 5*), Fornara et al. 2009<sup>d</sup> *FRI* (*FRIGIDA*), Stinchcombe et al. 2004<sup>e</sup> *LIF2* (*LHP1-INTERACTING FACTOR 2*), Latrasse et al. 2011<sup>f</sup> *MED12* (*MEDIATOR 12*), Imura et al. 2012<sup>g</sup> *COL5* (*CONSTANS-LIKE 5*), Hassidim et al. 2009<sup>h</sup> *MSI* (*MULTICOPY SUPPRESSOR OF IRA1*), Bouveret et al. 2006<sup>i</sup> *VIN3* (*VERNALIZATION INSENSITIVE 3*), Sung and Amasino 2004<sup>j</sup> *ZTL* (*ZEITLUPE*), Kim et al. 2007

**Supplementary Table 6:**
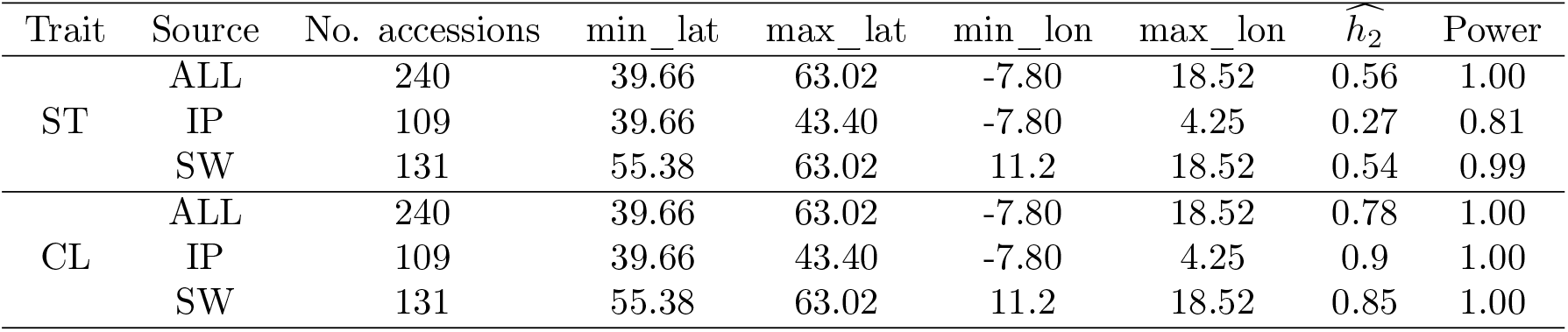
Geographic limits, pseudo-heritability and power estimation of the IP and SW subpopulation used for the analyses of stomata size (ST) and cauline leaf number (CL).

**Supplementary Table 7:** Summarized GWAS results from simulated data. True positives (TP), false positives (FP) and false discovery rate (FDR) are reported for 1,000 simulations per scenario with GWAS performed either in all 240 accessions (ALL) or only in the IP or SW subpopulation.

| Variance explained |  | TP |  |  | FP |  |  | FDR |  |  |
| --- | --- | --- | --- | --- | --- | --- | --- | --- | --- | --- |
|  |  | ALL | IP | SW | ALL | IP | SW | All | IP | SW |
| 20% | ALL | 964 | 264 | 390 | 225 | 73 | 146 | 0.19 | 0.22 | 0.27 |
|  | IP | 270 | 274 | 0 | 135 | 73 | 4 | 0.33 | 0.21 | 1 |
|  | SW | 65 | 0 | 420 | 187 | 0 | 156 | 0.74 | NA | 0.27 |

**Supplementary Table 8:** Overview of the sensitivity of the different simulation scenarios.

| Variance explained |  | Sensitivity |  |  |
| --- | --- | --- | --- | --- |
|  |  | ALL | IP | SW |
| 15% | ALL | 0.265 | 0.130 | 0.250 |
|  | IP | 0.050 | 0.000 | 0.230 |
|  | SW | 0.020 | 0.140 | 0.000 |
| 10% | ALL | 0.170 | 0.000 | 0.20 |
|  | IP | 0.010 | 0.000 | 0.020 |
|  | SW | 0.000 | 0.000 | 0.000 |
| 5% | ALL | 0.000 | 0.000 | 0.000 |
|  | IP | 0.000 | 0.000 | 0.000 |
|  | SW | 0.000 | 0.000 | 0.000 |

**Supplementary Table 9:** Overview of RNA expression data. This table shows the filters applied to the whole data. GWAS was performed on 2,483 genes.

| Filter | Sample size |
| --- | --- |
| Total RNA expression data | 24,175 |
| Nuclear genes | 23,021 |
| $\widehat{h}_2 > 0.5$ | 4,873 |
| $\widehat{h}_2 > 0.5$ & power > 0.9 | 2,483 |
| $\widehat{h}_2 > 0.5$ & power > 0.9 and not inflated | 1,982 |

## 10 Supplementary Files

Suppl. File 1 : Shared regions in the analyses of flowering time and associated candidate genes.

Suppl. File 2 : Summary of the number of associated SNPs and region for eGWAS on 2,237 genes.

Suppl. File 3 : List of 92 genes, which show different associations in the GWAS of the RNAseq data in the two subpopulation.

Suppl. File 4 : List of 110 genes, which show the same association in the RNAseq analysis in IP and SW.

Suppl. File 5 : List of 25 genes, which show an association in the RNAseq analysis in IP and SW subpopulation at the same region, but with different associated markers.

Suppl. File 6 : List of 118 genes, which show an association in the GWAS of the RNAseq data only in the IP subpopulation.

Suppl. File 7 : List of 64 genes, which show an association in the GWAS of the RNAseq data only in the SW subpopulation.

All files are accessible via https://github.com/arthurkorte/genetic_heterogeneity

